# Transcriptomic basis of within- and trans-generational predator-induced plasticity in the freshwater snail *Physa acuta*

**DOI:** 10.1101/2024.10.16.618730

**Authors:** Léo Dejeux, Nathanaëlle Saclier, Juliette Tariel-Adam, Maxime Hoareau, Tristan Lefébure, Lara Konecny, Sandrine Plénet, Emilien Luquet

## Abstract

Inducible defences in response to predation risk are a well-known example of adaptive phenotypic plasticity. Although inducible defences have been studied mainly within a generation (within-generational plasticity), there is now clear evidence that ancestral exposure to predation risk can influence the defences expressed by offspring, even if they have not been exposed themselves (transgenerational plasticity). The molecular mechanisms allowing the transmission of environmental information across generations are not well understood. In this study, we combined measures of antipredator responses (behavioural and morphological) with transcriptomic investigations across two generations in the freshwater snail *Physa acuta*. We hypothesised that both within- and transgenerational plasticity would induce phenotypic changes associated with differential gene expression. Our results confirmed within- and transgenerational plasticity: F1 snails respond to predator-cue exposure by increasing escape behaviour, reducing shell length, and developing thicker and slenderer shells, whereas F2 snails from exposed parents have longer and thicker shells with narrower apertures. Within- and transgenerational plasticity were accompanied by the differential expression of 112 genes (101 up- and 11 downregulated) and 23 differentially expressed genes (17 up- and 6 downregulated), respectively. Within- and transgenerational plasticity did not share common differentially expressed genes, but the associated molecular functions, involving metabolism and transcription regulation, were similar. These results suggest that predator-induced within-generational plasticity and transgenerational plasticity may result from different genomic pathways and may evolve independently.

## INTRODUCTION

Phenotypic plasticity is the ability of a genotype to produce alternative phenotypes under different environmental conditions (Pigliucci, 2005). Plastic responses are widespread across the phylogenetic spectrum (*e.g*. Galloway & Etterson, 2007; Luquet et al., 2011; Segers & Taborsky, 2011; Walsh et al., 2016) and are a major concern in evolutionary biology as they can be adaptive, i.e., increasing the fitness of organisms facing environmental changes (Pigliucci, 2005). Phenotypic plasticity classically refers to within-generational plasticity (WGP), meaning that it occurs within one generation. Recently, plasticity has also been found to occur across generations (transgenerational plasticity [TGP]), where the phenotype of an organism is influenced by the environment experienced by previous generations (Bell & Hellmann, 2019). Interest in TGP has grown rapidly in recent years with evidence that TGP is at least as widespread as WGP (Jablonka & Raz, 2009; Tariel, Plénet, et al., 2020b). Although there are numerous examples of TGP generating seemingly adaptive offspring phenotypes, its adaptive role is expected to be more challenging than that of WGP because of patterns of environmental variation across generations (Colicchio & Herman, 2020). Indeed, the selection of a strong temporal autocorrelation between environments of distinct generations (Ezard et al., 2014; Kuijper & Hoyle, 2015; Prizak et al., 2014) and a transmission system that allows the induction of adaptive phenotypes across generations is required (Fallet et al., 2020). The inheritance mechanisms appear diverse, ranging from epigenetic changes to parental transmission of cytoplasmic elements (small RNAs, nutrients, hormones, proteins), parental care and cultural transmission (Bell & Hellmann, 2019; Danchin et al., 2011).

Regardless of the transmission mechanism, the influence of the parental environment on the offspring phenotype should be linked to changes in gene expression, at least in some specific tissues and developmental stages. Surprisingly, the transgenerational plastic response at the molecular level has been overlooked. This is indeed a necessary first step to unravel the molecular mechanisms of plasticity in general and their transgenerational inheritance. The most compelling explorations of changes in gene expression profiles in response to both developmental (WGP) and parental (TGP) environments have been conducted by Hales et al. (2017) on *Daphnia ambigua* and (Stein et al. (2018) on the three-spined stickleback (*Gasterosteus aculeatus*). These studies compared gene expression between individuals that had been exposed to predator cues relative to nonexposed individuals within a generation (WGP genes) and among their offspring (from non-exposed and exposed parents; TGP genes). Stein et al. (2018) reported that the expression of the same group of genes changed their expression in offspring regardless of wether predation risk was experienced by the offspring (TGP), the father (WGP), or both. This suggests that a core set of genes is consistently affected by predation risk regardless of who experiences it. Conversely, Hales et al. (2017) reported that changes in the expression of genes that respond to TGP differed depending on whether exposure to predation was experienced by parents or grandparents. They also revealed that very few genes were common to both WGP and TGP. Notably, Clark et al. (2019) also reported that the TGP gene expression profiles of larvae of sea urchins (*Psammechinus miliaris*) spawned at low pH from preacclimated adults differed from those of larvae produced by adults exposed to ambient pH, but the authors did not explore the WGP genes. These few results highlight the necessity of investigating WGP and TGP together at the molecular level. In addition to the inheritance mechanisms allowing organisms to retain information about past environmental conditions to adjust their future phenotype, we need to know whether the two forms of plasticity are distinct at the molecular level and potentially free to evolve independently of one another (Bell & Stein, 2017).

Predation risk is a major selection pressure for most species and results in the evolution of defences in prey (Kikuchi et al., 2023). As predation risk often varies over time and space, it favours the evolution of phenotypic plasticity in defences (Reger et al., 2018; Tariel, Plénet, et al., 2020b; Viney & Diaz, 2012). Indeed, although some defences against predators are constitutive, plastic defences are only induced when prey detect some predator cues in their environment (*e.g*. visual cues, chemical cues such as predator odours, or alarm signals released by conspecifics). These induced defences are classic examples of adaptive WGP: prey can increase their fitness by producing defences only when predators are present. The links between prey-predator interactions and WGP have long been studied (*e.g.* Tariel et al., 2020b; Tollrian, 1995). Plastic defences can also be induced in offspring after their parents or more distant ancestors have detected predator cues. This transgenerational induction of defences was first highlighted in Daphnia morphology by Agrawal et al. (1999) and has since been repeatedly shown in multiple species and defences (review in MacLeod et al., 2022; Tariel, Plénet, et al., 2020b).

Consequently, this work aims to elucidate the molecular mechanisms underlying the expression of WGP and TGP in defences in the context of predator-prey interactions. We used *Physa acuta* (also referred to as *Physella acuta*), a freshwater snail known for its within- and transgenerational antipredator defences, such as escape behaviour e.g.,snails crawl out of the water when they or their parents detect predator cues (Luquet & Tariel, 2016; Tariel, Luquet, et al., 2020; Tariel, Plénet, et al., 2020a; Turner et al., 1999), changes in shell size and shape, increases in shell thickness and overall increases in shell-crush resistance (*e.g.*, shell are harder to handle or to crush by predators (Auld & Relyea, 2011; Beaty et al., 2016; DeWitt, 1998; Tariel, Plénet, et al., 2020c; Tariel-Adam et al., 2023). We combined a two-generation experiment to generate predatorinduced WGP and then TGP of defences (only in a nonexposed environment) with a transcriptomic investigation. We used transcriptome sequencing and assembly to investigate transcriptome mantle skirt gene expression specifically (the mantle skirt is the tissue that synthesises the shell). We expected that gene expression would be affected by current (WGP) and parental (TGP) exposure to predator cues. More specifically, as WGP and TGP induce similar phenotypic responses (*e.g.* increases in shell thickness), we expected the two processes to share the same core set of genes.

## MATERIALS & METHODS

### Animal collection, rearing conditions and experimental design

Adult *P. acuta* snails were collected from a wild population in the lentic backwater of the Rhône River (Lyon, France, 45° 48’6"N, 4° 55’33"E) in September 2018. Individuals from this population have been repeatedly used in previous studies on predator-induced plasticity (Luquet & Tariel, 2016) and are naturally exposed to predation by the crayfish *Faxonius limosus*. The experiment was conducted in our laboratory in a temperaturecontrolled chamber at 25 °C with a photoperiod of 12/12 h. The wild snails constituting the F0 generation were pooled in a 10 L plastic box filled with dechlorinated tap water (control water hereafter) and interbred overnight. They were then isolated in 80 mL plastic boxes (4.5 × 6 cm) filled with tap control (i.e., without predator cues) water to lay eggs for 3 days. Eggs within boxes constitute F1 matrilines. Eggs hatched seven days later, and siblings subsequently grew together for an additional ten days in water without predator cues. Then, 10 siblings of 20 F1 matrilines were randomly selected and split into two developmental environments: five siblings in water without predator cues (nonexposure treatment) and five siblings in water with predator cues (exposure treatment). F1 siblings individually developed for the next 31 days in 80 mL plastic boxes. On this date, one F1 snail per matriline and treatment was frozen at −80 °C for molecular analysis and all remaining F1 snails were measured (see below).

To generate the F2 generation, we pooled F1 snails from the same treatment in an aquarium filled with control water and allowed them to copulate for 24 h. As previously described, F1 snails were then isolated in 80 mL boxes and laid eggs for 3 days to produce F2 matrilines. F2 snails from the two parental environments (nonexposed and exposed to predator cues) were reared only in control water to investigate the effect of the parental environment (TGP) alone, i.e., without the interaction between parental and developmental environments. Our previous study revealed that TGP is stronger in a mismatch situation (i.e., when F2 from exposed parents are not exposed (Luquet & Tariel, 2016)). We followed the same protocol steps as previously described for F1 snails for F2 generation, except that eight siblings per matriline were reared and measurements were made at an older age because of slower growth. For F1, one F2 snail per matriline and treatment was frozen for molecular analysis, and the others were measured.

Snails were fed *ad libitum* with boiled and mixed lettuce. The water (according to the treatment) and food were renewed twice a week. The water with predator cues (i.e., exposed treatment) was the water in which *F. limosus* crayfish were raised (one crayfish/5 L), for 3 days and were subsequently fed with snails. Additionally, to ensure the presence of fresh alarm cues, we smashed several *P. acuta* adult snails in predator-cue water (one snail/5 L) for 1 h before use.

### Behavioural and morphological responses

In this study, to check that exposure to predator cues triggered WGP and TGP, escape behaviour and shell morphology were measured in F1 (nonexposed snails = 76, exposed snails = 73) and F2 (snails from nonexposed parents = 89, snails from exposed parents = 101).Some mortality events led to uneven sample sizes, but the mortality rates remained consistent across families (see Table S1). Crawling out of the water (positioning themselves above the water surface) allows snails to escape benthic predators such as crayfish (DeWitt, 1998; Tariel, Plénet, et al., 2020a). Shorter and narrower shell and aperture dimensions and thicker shells are adaptive antipredator responses (Auld & Relyea, 2011). These phenotypic traits have consistently been implicated in the transgenerational responses of *P. acuta* to predator cues (Beaty et al., 2016; Luquet & Tariel, 2016).

*Escape behaviour:* One week before the measurements and four hours after the water renewal, we recorded the position above or below the water surface of each snail in their rearing boxes. Consequently, the behaviour of F1 snails was recorded in water without or with predator cues according to their respective treatments, whereas that of F2 snails was recorded only in water without predator cues.

*Shell thickness, shell length and shape*: Shell thickness was directly measured with an electronic calliper at the nearest 0.01 mm at the edge of the aperture. A photograph of each snail (aperture upwards) was subsequently taken with an Olympus SC50 camera installed on an Olympus SZX9 binocular and an Olympus DF PLAPO 1X-2 objective at a 8× magnification. Snail shell length, shell width, aperture length and aperture width were measured using ImageJ (Schneider et al., 2012). To estimate potential changes in shell shape according to treatment, we used geometric methods (Gustafson et al., 2014). For each snail shell, we digitised 19 landmarks (Fig. 1) using ImageJ software. Snails that lacked homologous landmarks because of broken shells (typically broken apertures and apexes) or small sizes (lack of the third shell spire) were removed from the analyses (nonexposed F1 snails = 76, exposed F1 snails = 72, F2 snails from nonexposed parents = 89, F2 snails from exposed parents = 99). To compare shell shapes according to treatment, we scaled the shapes using the size metric defined below (Eqn. 1). The rotational components were removed by translating all shapes so that the centroids lie at the origin. Rotational components were removed by rotating all the shapes until the Procrustes distance was minimised, with the first shell fixed as a reference.

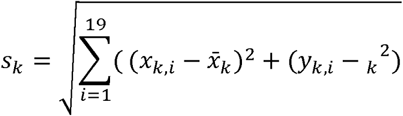

**Equation 1**: Procustes equation used to scale the shapes using the size metric , which is the size of the -th shell; and are the x and y components of the -th landmark; ( ) is the centroid of the -th shell.

**Figure 1:**
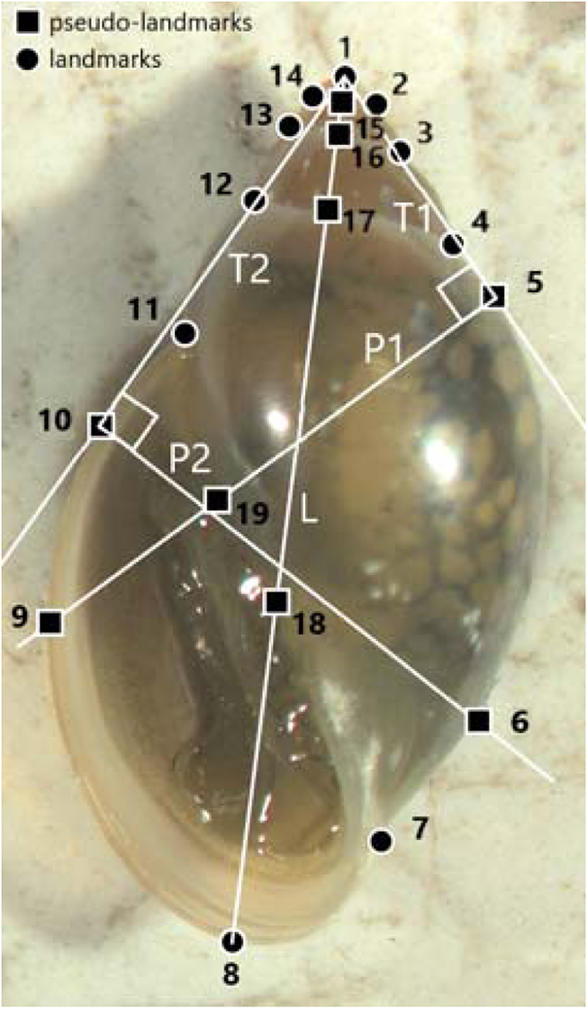
Locations of the different landmarks (circle symbols) and pseudo-landmarks (positioned using a geometric method, square symbols) on *P. acuta* shell. Pseudo-landmarks were created using a geometric method: first, two tangents (T1 and T2) were drawn from landmark 1 (apex). T1 touches the curve of the habitation spire (5) and T2 touches the external edge of the aperture (10). From the two points of tangency (points 5 and 10), two lines perpendicular to the tangents were traced (P1 and P2). The intersections with the columellar edge of the aperture, the external edge of the aperture and the curve of the habitation spire give additional pseudo-landmarks (6, 9, 18 and 19). The length of the shell L (straight line passing by 1 and 8) gives pseudo-landmark 18 at the intersection with the columellar edge of the aperture).

### Statistical analyses

The same statistical analyses were separately performed to test for WGP on F1 snails and TGP on F2 snails. The treatment factor (i.e., nonexposed or exposed to predator cues) was the developmental environment for WGP (F1), whereas it was the parental environment for TGP (F2).

*Escape behaviour:* The effects of predator cues on escape behaviour (i.e. snail position above/on or below the water surface) were analysed using generalised linear mixed models (GLMMs) assuming a binomial distribution (logit link function). The treatment was a fixed effect, and matriline was a random intercept effect. We tested the significance of fixed and random effects with likelihood ratio tests.

*Shell thickness and shell length:* To analyse the effects of predator cues on shell length and shell thickness, we performed linear mixed models with treatment as a fixed effect and matriline as a random intercept. We used restricted maximum likelihood estimation and Kenward and Roger’s approximation for degrees of freedom. We tested the significance of fixed effects with type II F-tests (Kuznetsova et al., 2017) and the significance of random effects with likelihood ratio tests.

Linear mixed models and generalised linear mixed models were generated using the lme4 package (Bates et al., 2014) in R software (version R-4.4.1 for Windows, (RStudio Team, 2020).

*Shell shape:* The effects of predator cues on shell shape were analysed using principal component analysis (PCA) and visualised with the 2D landmark relative warp analysis functionality. All the standardised shapes obtained via the Procrustes method were analysed using PAST software with a Student test (Hammer et al., 2001).

**RNA Extraction and Sequencing** The mantle skirt was dissected for 5 nonexposed F1 snails and 5 exposed F1 snails, and for 10 F2 snails from nonexposed parents and 10 F2 snails from exposed parents. Total RNA from the mantle was extracted by adding TRI Reagent (Molecular Research Center MRC, TR118) to the samples and homogenising the tissues on ice with pistons and an agitator (Motor Mixer Clearline ref.045011 Dutscher). All remaining steps were carried out according to the manufacturer’s protocols (Molecular Research Center MRC, TR118). Then, the RNA was treated with Turbo DNase enzyme (Turbo DNA free kit, Invitrogen, AM1907) and assayed by fluorescence using a Qubit fluorometer (Qubit® RNA HS Assay Kits Molecular Probes, Invitrogen, Q32855). Library construction was carried out using an NEBNext Ultra II RNA Library Prep Kit for Illumina (Biolabs New England, NEB E7770, E7490, E7600). Libraries were randomly split into three groups and sequenced on three lanes with an Illumina® HiSeq4000^™^ machine on the GenomEast platform hosted at the IGBMC (GenomEast Platform – Institut de Génétique et de la Biologie Moléculaire et Cellulaire IGBMC, Illkirch, France), resulting in 50 bp long single-end reads.

### Differential gene expression analysis

FASTQC (http://www.bioinformatics.babraham.ac.uk/projects/fastqc/)) was used to assess the quality of the reads. We used a reference genome ORF *P. acuta* available on NCBI (Isolate: Inbredline_101_S28. RefSeq: GCF_028476545.12). Gene expression levels were assessed on annotated transcripts using Kallisto (-l 50 -s 10 options; Bray et al., 2016). We aggregated effective counts at the gene level and then applied rounding to the nearest whole number to convert noninteger values into integer counts. We also applied a minimum count filter of 1 (cpm) where we removed all the genes with reads below 1 (i.e. removing 5634 genes from the total of 46,965). The rounded effective counts were used as inputs for differential gene expression using the DESeq2 package (Love et al., 2014), which performs differential gene expression analysis using negative binomial distribution and shrinkage estimation for dispersions (“apeglm” type shrinkage, Zhu et al. 2019), and outputs expression fold changes to improve the interpretability of the estimates. We specified nonexposed or exposed treatment in the model (design formula: “design = ∼treatment”) within each generation independently, meaning that we compared F1 snails from nonexposed and exposed treatments, and F2 snails from nonexposed and exposed parents. First, the Wald test (DESeq2 package) was used to test the significance of differentially expressed genes (DEGs) in all samples between the nonexposure and exposure groups. Second, we checked for any structures in the normalised count table by performing a principal component analysis (PCA) of the 500 most DEGs from DESeq2 analysis (pval < 0.05) based on Log2FoldChange (|LFC|, i.e., the logarithm base 2 of the fold change in gene expression between two experimental conditions, providing a measure of both the magnitude and direction of differential expression). Third, we filtered the significant DEGs by an |LFC| greater than 1 in each generation. Consequently, only differential expression levels with adjusted p-values < 0.05 and |LFC| > 1 were considered thereafter. We subsequently investigated whether the DEGs in F1 were also differentially expressed in F2 by comparing specific gene IDs across the datasets from the two generations. Finally, as the quality of annotations of the reference genome did not allow complex functional analyses, we determined the potential gene functions and the associated protein families of all significant DEGs using the scan procedure of InterPro, a tool provided by the European Bioinformatics Institute from the European Molecular Biology Laboratory (EMBL-EBI; (Blum et al., 2024; Jones et al., 2014). InterPro integrates signatures from 13 member databases to classify proteins into families, to predict functional domains, and to identify key sites, enabling comprehensive functional annotation.

## RESULTS

### Predator-induced behaviour and shell morphology

#### Within-generational Plasticity

In the F1 generation, the proportion of crawling-out behaviour increased significantly in snails exposed to predator cues (+85.25%, F[1,19.43] = 321.82, p < 0.001; Fig. 2A). Shell length significantly decreased in snails exposed to predator cues (−6.3%, F[1,128.71] = 15.46, p < 0.001; Fig. 2B). Shell thickness significantly increased in exposed snails (+18.8%, F[1,129.58] = 4.02, p = 0.047; Fig. 2C). The matriline random effect was significant for shell length (χ^²^[1] = 19.022, p < 0.001) and nonsignificant for behaviour and shell thickness (χ^²^[1] = 0.690, p = 0.406 and χ^²^[1] = 2.6116, p = 0.106, respectively).

**Figure 2:**
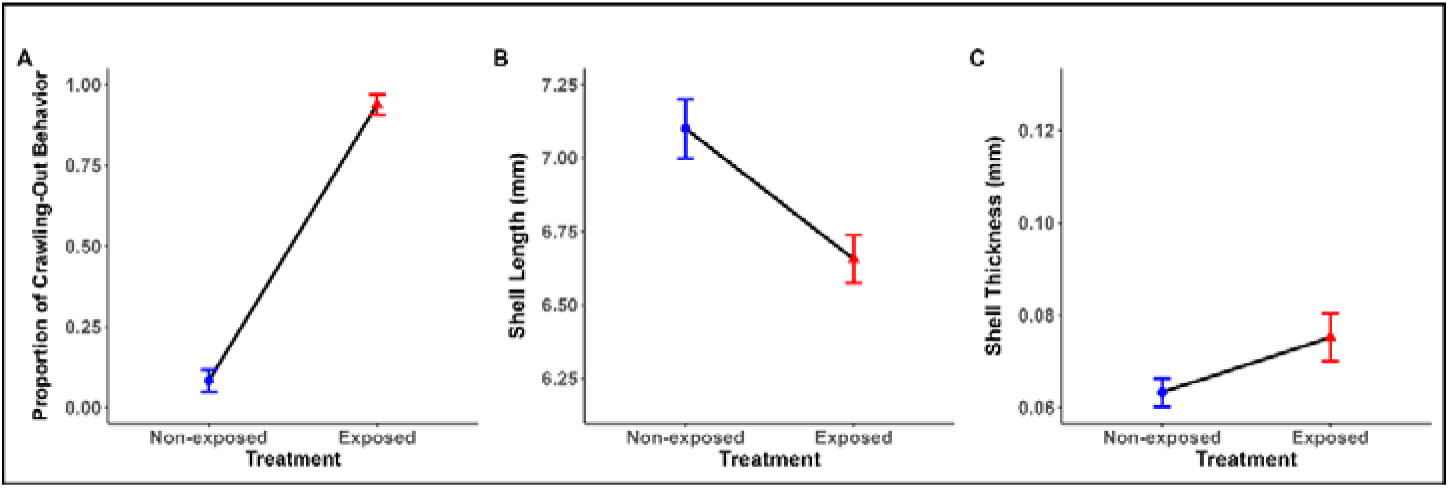
Effects of developmental exposure to predator cues (WGP) on crawling-out behaviour, shell length and shell thickness in the F1 generation. A: Proportion of crawling-out behaviour; B: mean shell length (mm) ; C: mean shell thickness (mm). Error bars represent SE. Nonexposed and exposed to predator-cue treatments are symbolised with blue circles and red triangles, respectively.

Exposure to predator cues induced changes in shell shape. The first two axes of the PCA accounted for 59.2% of the shell shape variation (Fig. 3), and PCA axes 1–4 (74.2% cumulative shape variation) were the only axes accounted for more than 5% of the shape variation. PC1 showed a change in the columellar area (close to landmarks 18–19) from shells with relatively wide apertures (dilated columellar area; Fig. 3C) to shells with relatively narrow apertures (compressed columellar area; Fig. 3D). PC2 changed in the slenderness of the shell from shells with relatively short spires (compressed apex; Fig. 3B) and relatively wide apertures (dilated aperture) to shells with relatively short spires (dilated apex) and narrow aperture widths (compressed aperture; Fig.3A).

**Figure 3:**
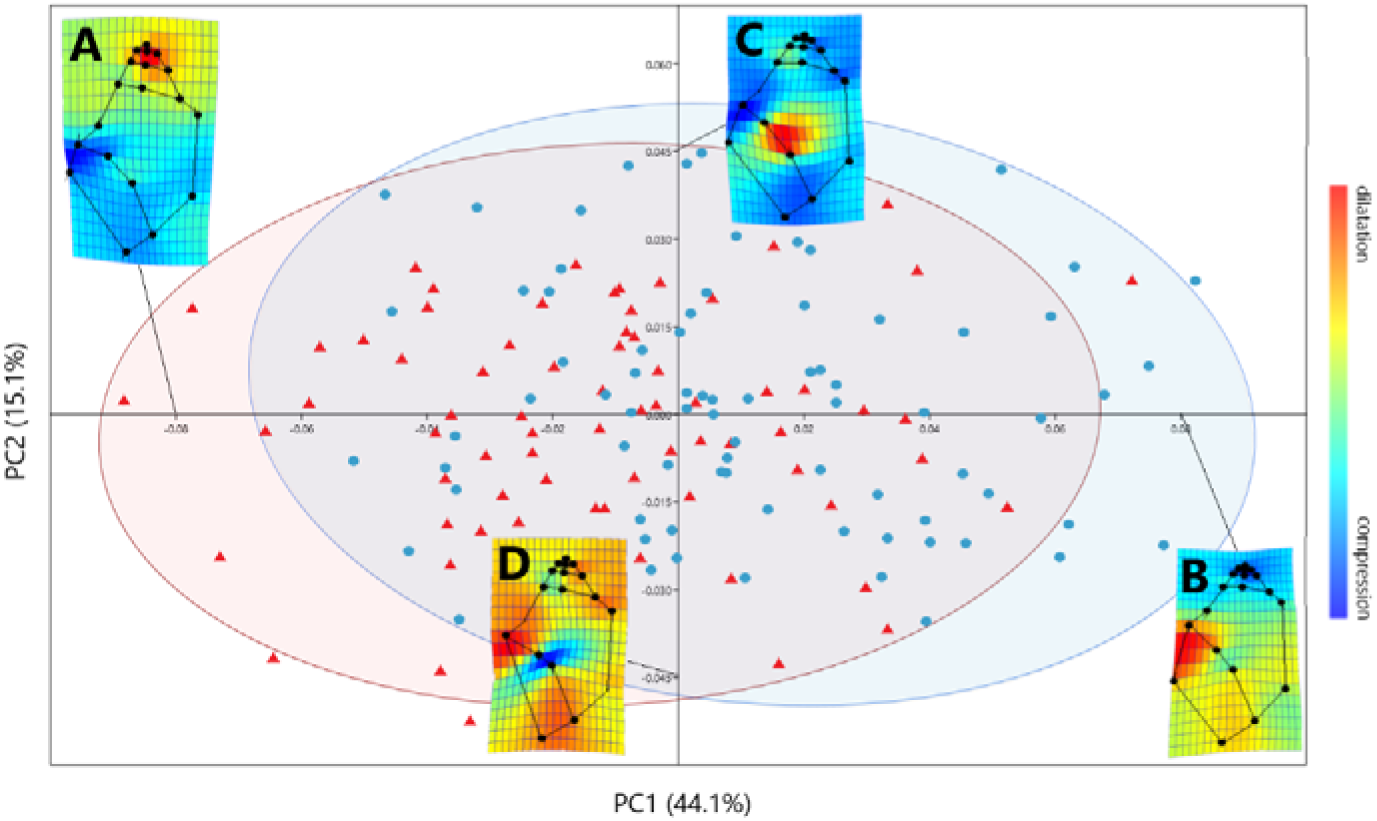
Shape variation of nonexposed and exposed snails from the F1 generation (blue circle and red triangle symbols, respectively. Panels A, B, C, and D represent deformation grids with shell shapes on the extreme ends of the PCA axes. The 95% confidence ellipses are represented for nonexposed and exposed to predator-cue treatments. The dilation□compression gradient bar shows the deformation of the shell landmarks relative to the points of an average shell (which would be located at the centre of the PCA).

Compared to exposed snails, nonexposed snails presented the same range of deformation in terms of the columellar area (PC2, t-test: t = 1.07, df = 144, p = 0.28). However, the shells of snails exposed to predator cues were slenderer, with longer spires and narrower aperture widths than those of nonexposed snails (PC1, t-test: t = 4.803, df = 144, p < 0.001). This elongation was associated with the dilatation of the last three spires and a narrow aperture at the base of the external edge. In contrast, the shells of nonexposed snails had a wider aperture at the base of the external edge and smaller spires.

#### Transgenerational plasticity

In the F2 generation, the proportion of crawled snails did not differ between snails from nonexposed and those from exposed parents (F[1,20.31] = 3.929, p = 0.061; Fig. 4A). Snails from parents exposed to predator cues were significantly longer and had thicker shells than those from nonexposed parents (+4.9%, F[1,170.87] = 7.55, p = 0.006 and +16.9%, F[1,174.11] = 9.858, p = 0.002, respectively; Fig. 4B-C). The matriline random effect was significant for shell length (χ²[1] = 31.685, p < 0.001) and nonsignificant for behaviour and shell thickness (χ²[1] = 0.022, p = 0.882 and χ²[1] = 1.882, p = 0.17, respectively).

**Figure 4:**
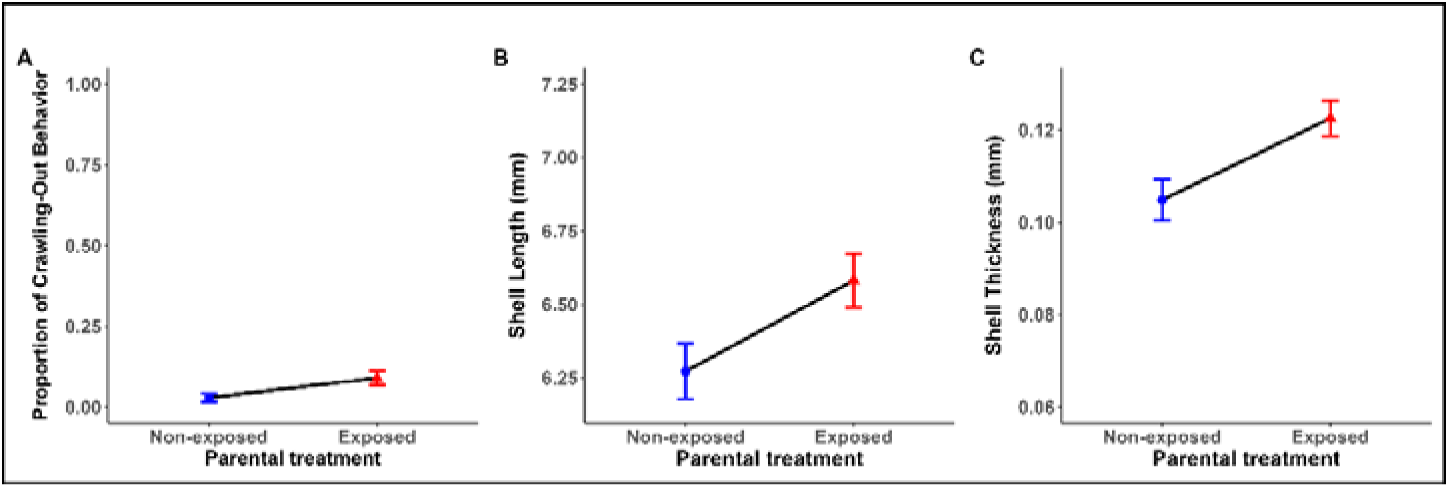
Effects of parental exposure (TGP) on crawling-out behaviour, shell length and shell thickness in the F2 generation. A: Proportion of crawling-out behaviour; B: mean shell length (mm) ; C: mean shell thickness (mm). Error bars represent SE. Non-exposed and exposed to predator cues are symbolised with blue circles and red triangles, respectively.

Parental exposure to predator cues induced changes in shell shape. The first two axes of the PCA accounted for 63.8% of the shell shape variation, and the cumulative percentage accounting for PCA axes 1–4 was 80.14% (Fig. 5). PC1 displayed a change in shell aperture width; the shells with high PC1 scores (Fig. 5C) showed a dilatation of the shell where the outer lip meets the parietal wall and a small compression of the columellar area (i.e., a wider aperture). Shells with low PC1 scores (Fig. 5D) presented a narrower aperture, a consequence of a dilated columellar area but compression of the parietal wall and basal lip in comparison to shells with high PC1 scores. PC2 showed a change in shell aperture width but also in shell length, ranging from shells (Fig. 5C) with a dilated outer lip and compression of the apex (which tends to reduce spire length, i.e., overall shell length) to shells (Fig. 5D) showing elongated spires from the dilated apex, with a narrower aperture due to outer lip compression.

**Figure 5:**
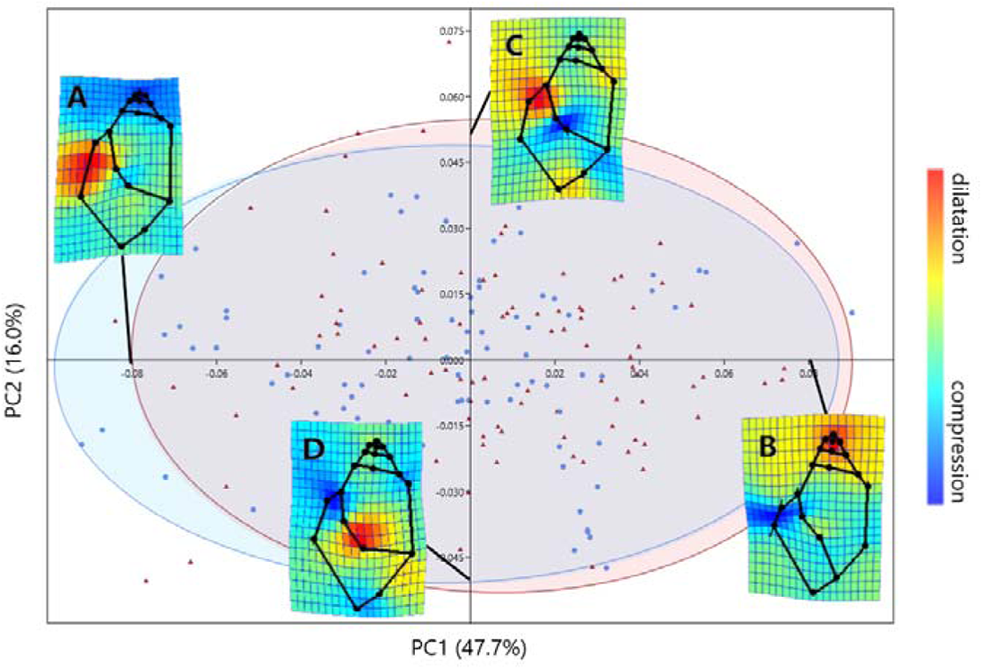
Shape variation of F2 snails from nonexposed and exposed parents (blue circles and red triangles symbols, respectively). Panels A, B, C, and D represent deformation grids with shell shapes on the extreme ends of the PCA axes. The 95% confidence ellipses are represented for nonexposed and exposed to predator-cue treatments. The dilation□compression gradient bar shows the deformation of the shell landmarks relative to the points of an average shell (which would be located at the centre of the PCA).

Snails from predator-exposed parents had a dilatation of the shell in the columellar area accompanied by a small compression of the parietal wall (PC1: t = 2.067, p = 0.04). Conversely, there was no difference in deformation of the outer lip or dilatation or compression of the apex between snails from exposed and nonexposed parental environments (PC2: t = 0.59, p = 0.55).

### Differential gene expression induced by predator-cues

A principal component analysis (PCA) was performed to explore the clustering of read counts on the first 500 most DEGs. The first and second principal components (PCs 1 and 2) explained 18% and 11% of the variance respectively (Fig. 6, PCA loadings available in Fig. S1). The plot did not show clustering between the nonexposed and exposed treatments or between the F1 and F2 generations (Fig. 6).In the F1 generation, differential expression analysis revealed a total of 112 DEGs between nonexposed and exposed snails (101 upregulated genes and 11 downregulated gene; Fig. S2). In the F2 generation, the differential expression analysis revealed a total of 23 DEGs between snails from nonexposed and exposed parents (17 upregulated and 6 downregulated gene; Fig. S2). A comparison of DEGs in F1 and F2 revealed that they did not overlap (Fig. 7; Fig. S3).

**Figure 6:**
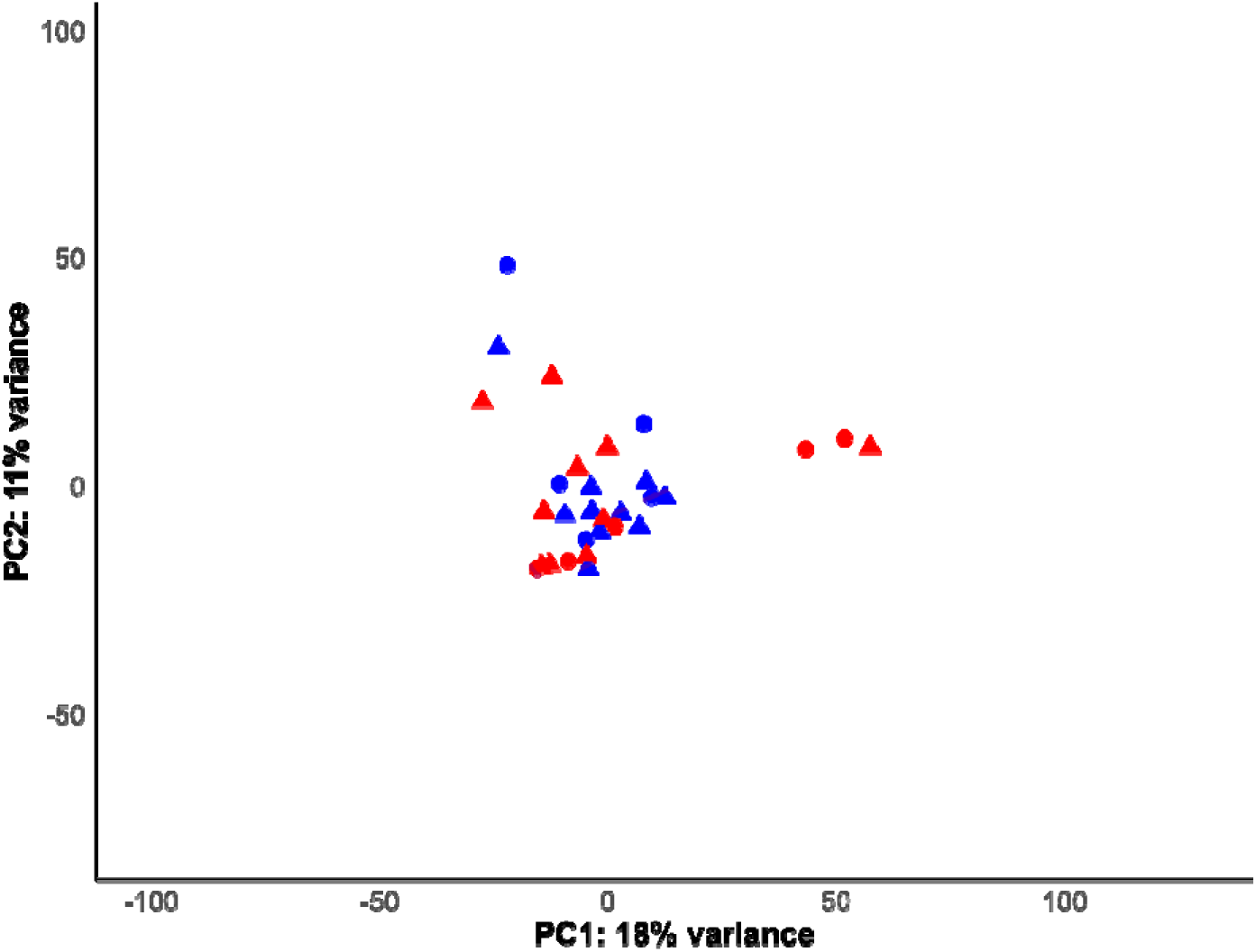
PCA plot of the 30 transcriptome samples of *P. acuta* snails showing the divergence in differential expression among the first 500 most differentially expressed genes (selected by greatest |LFC|). Nonexposed and exposed to predator cue individuals are represented in blue and red, respectively. F1 individuals are represented by circles and plain ellipses, and F2 individuals are represented by triangles and dotted ellipses.

**Figure 7:**
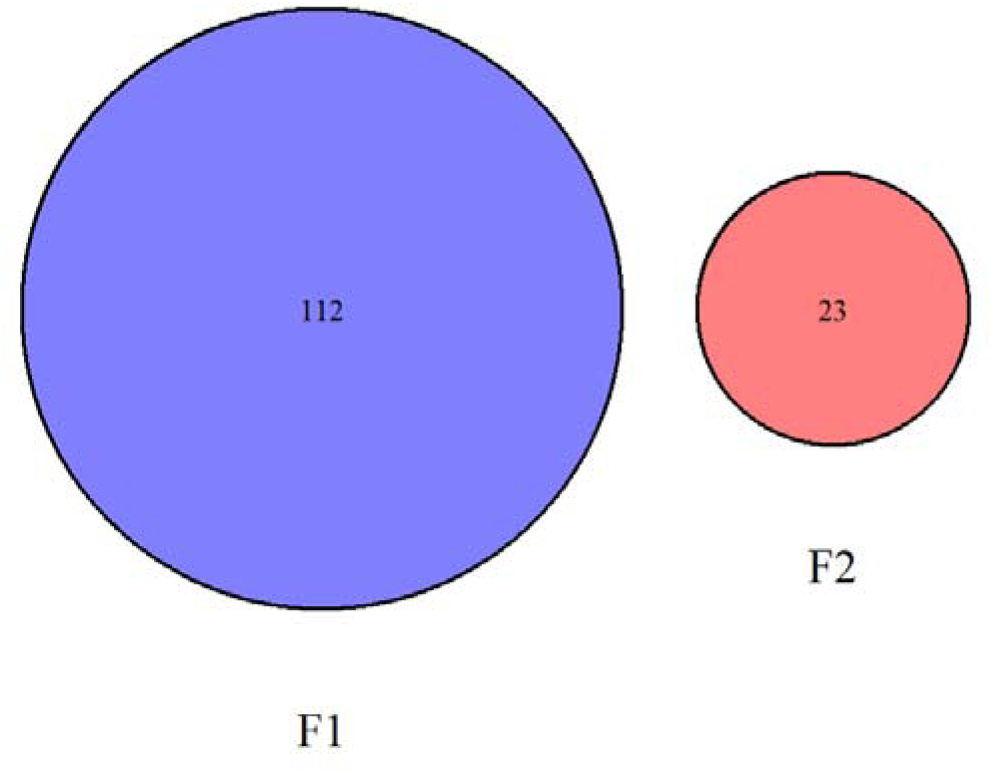
Venn diagram (obtained with the DESeq2 package) illustrating the number and overlap of differentially expressed genes between the F1 and F2 groups. Genes specific to each group are shown in individual circles; the intersection represents the shared genes.

The DEGs in F1 were involved mainly in metabolism, signal transduction, extracellular matrix organisation and protein and transcriptional regulation (Fig. 8, Tables S2 and S3 for further details). The DEGs in F2 were involved mainly in metabolism and transcriptional regulation, including DNA methylation regulation (Fig. 8, Tables S2 and S3 for further details). In both F1 and F2, some DEGs were associated with calcium signalling (Fig. 8, Tables S2 and S3 for further details), and the majority of DEGs were genes with unknown functions.

**Figure 8:**
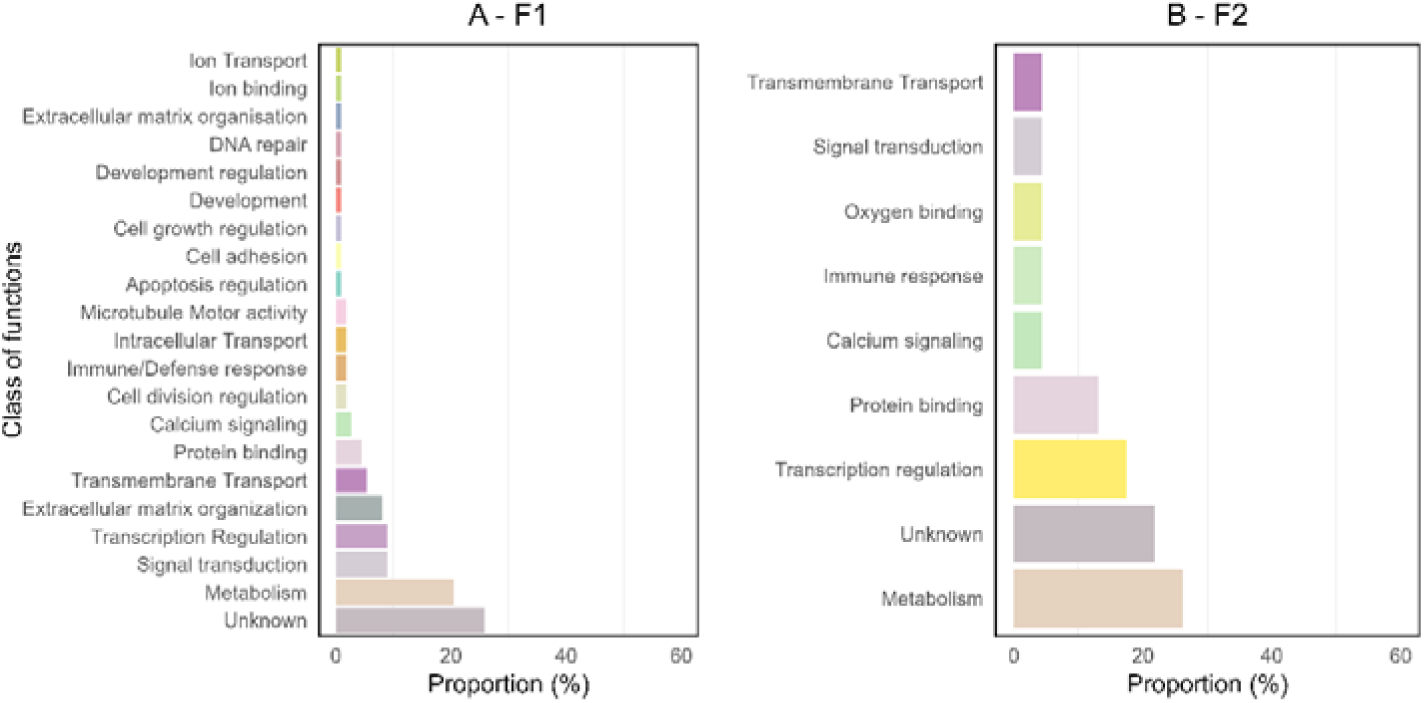
Proportions of classes of functions of the differentially expressed genes in F1 (A) and F2 (B) based on the InterPro EMBL-EBI database (Jones et al., 2014; Blum et al., 2024).

## DISCUSSION

This work aimed to elucidate the molecular mechanisms underlying the expression of within- and trans-generational plasticity of antipredator defences in *P. acuta*. We expected that gene expression would be affected by developmental (WGP) and parental (TGP) exposure to predator cues and that WGP and TGP would share the same core set of genes. Exposure to predator cues resulted in changes in escape behaviour, shell growth, shell thickness and shell shape. The parental exposure to predator cues (TGP) also influenced the offspring phenotype, particularly morphological traits as previously shown in this system (*e.g*., Luquet & Tariel 2016). Accordingly, DEGs were detected between nonexposed and exposed snails in F1 (WGP) and between F2 snails according to parental exposure to predator cues (TGP). However, the number of DEGs detected for TGP was lower than that detected for WGP. Finally, our results revealed that DEGs associated with WGP and TGP did not overlap.

### Phenotypic changes and gene expression induced by within-generational exposure to predator cues

As expected, snails exposed to predator cues crawled more out of the water, had thicker shells, were smaller and developed slenderer shells with longer spires and narrower aperture widths than nonexposed snails did. These behavioural and morphological responses represent defences and are supposed to be adaptive as they are consistent with escape behaviour (Tariel, Luquet, et al., 2020) and an increase in shell crush resistance (Tariel, Plénet, et al., 2020a, 2020c), increasing snail survival in the face of crayfish predation (Auld & Relyea, 2011). The decreased shell size observed in exposed snails may result from physiological stress caused by exposure to predator cues, including costs to produce antipredator defences (*e.g*., crawling-out behaviour occurs at the expense of foraging; DeWitt, 1998).

These predator-induced WGP changes were associated with a distinct pattern of gene expression with 112 DEGs (101 upregulated and 11 downregulated genes). Overall, this pattern is consistent with several studies that explored the changes in gene expression induced by the environment (*e.g*., Aubin-Horth & Renn, 2009; Herman & Sultan, 2011). In the context of predator=prey interactions, the main knowledge about the transcriptomic responses of prey to predators comes from studies on Daphnia species. For example, Hales et al. (2017) in *Daphnia ambigua* and Rozenberg et al. (2015) in *Daphnia pulex* reported that antipredator defences, mainly those related to morphological traits, were associated with 48 and 230 DEGs respectively. In contrast, no DEGs were identified in *Daphnia galeata,* a species known to adjust life history traits (such as reproduction) but not morphological traits in response to predator cues. Similarly, Orsini et al. (2018) revealed no changes in gene expression associated with short-term exposure to fish kairomones in *Daphnia magna*. These contrasting results suggest that the patterns of gene expression may depend on several factors such as the traits considered and the strength of predator signals (*e.g*. cue concentration and duration of exposure).

The functional analysis of DEGs in snails exposed to predator cues suggested that these genes are linked mainly to metabolic functions and transcription regulation, both of which are closely associated with stress responses. Notably, 26% of the DEGs had unknown functions. A substantial portion of these genes are involved in generic metabolic processes (e.g., complex I intermediate-associated protein 30, which plays a role in respiratory metabolism; Walker et al., 1992). These functions are essential for maintaining cellular stability and enabling adaptation to fluctuating environmental conditions. However, some genes are implicated in more stress-specific responses that may be directly triggered by predator exposure (*e.g*., involvement in the TOR protein complex, which is well known to influence the response to abiotic stressors; Blackwell et al., 2019; Katewa & Kapahi, 2011). Similarly, some DEGs were linked to calcium ion metabolism (*e.g*., by coding for the annexin and peptidase C2 protein families; Khorchid & Ikura, 2002) and could be involved in shell synthesis. Calcium is the main component of shell gastropods (Bukowski & Auld, 2014), and modulations of general calcium pathways could influence the expression of antipredator defences, such as increases in shell thickness and changes in shell shape, as observed in our study. Our findings align with previous research investigating the functions of DEGs in predator cue environments and showing that genes are associated with defensive structures, metabolism and transcription regulation. Indeed, Hales et al. (2017) and Rozenberg et al. (2015) reported that the gene functions of *D. ambigua* and *D. pulex* mostly explained the observed defensive phenotypic changes (*e.g*., the expression of proteins involved in the formation of protective structures and in cuticle strengthening), as well as the metabolic pathways involved in resource allocation and stress responses. Similar results have been reported in larvae of the damselfly *Enallagma cyathigerum* and in the marine copepod *Calanus finmarchicus,* where predation risk activated genes encoding for stress proteins, reduced antioxidant defence, and altered lipid metabolism (Skottene et al., 2020; Slos & Stoks, 2008). Although the relationship between the functions of DEGs and phenotypic traits is highly speculative in nonmodel organisms, our study may highlight some interesting molecular pathways underlying predator-induced WGP in *P. acuta* that would be interesting to study in further detail.

### Phenotypic changes and gene expression induced by transgenerational exposure to predator cues

Our results confirmed predator-induced TGP. The parental exposure to predator cues significantly influenced the offspring phenotype, particularly the morphological traits. F2 snails from exposed parents had thicker and longer shells with narrower apertures than snails from nonexposed parents while they had similar antipredator behaviour. These transgenerational effects are consistent with previous studies on this system which demonstrated that parental environments shape the antipredator traits of offspring (Beaty et al., 2016; Tariel, Plénet, et al., 2020c). A thicker shell and a larger size increase the shell-crush resistance, and a narrower aperture limits body snail extirpation by crayfish (Auld & Relyea, 2011; DeWitt, 1998). The longer shell of F2 snails from exposed parents was opposed to the shorter shell of exposed snails in F1. This contrasting pattern aligns with our previous study (Tariel, Plénet, et al., 2020c), suggesting that offspring of exposed parents, having received information about predation risk at an early developmental stage, activate pathways that enable simultaneous investment in shell length to reach refuge size (Auld & Houser, 2015) and shell thickness for increased crush-resistance (Beaty et al., 2016; Tariel-Adam et al., 2023). The antipredator behaviour was similar between snails from nonexposed and exposed parents, which is in line with some contrasting results reported in other studies (Tariel, Plénet, et al., 2020a, 2020c). Overall, the expression of antipredator traits by offspring from exposed parents, even if they are not themselves exposed to predator cues, may allow them to anticipate future predation risk and increase their survival (MacLeod et al., 2022; Tariel, Plénet, et al., 2020b).

At the molecular level, predator-induced TGP was associated with 23 DEGs (17 upregulated and 6 downregulated) according to the parental environment. The low number of DEGs in the F2 generation contrasts with studies on factors other than predation (*e.g.*, pH in Clark et al., 2019; temperature in Ledón-Rettig, 2023) and with parental exposure to predator cues (Hales et al., 2017; Mommer & Bell, 2014; Stein et al., 2018). For example, Stein et al., (2018) found 322 DEGs in three-spined stickleback *G. aculeatus* offspring when predation risk was experienced by the father. In *D. ambigua*, individuals from predator-exposed parents and grand-parents presented 233 and 170 DEGs, respectively, compared with individuals from nonexposed parents (Hales et al., 2017). Although this contrast may result from a less stringent method of DEG selection (no LFC filtering, *e.g*. Stein et al. 2018), it also raises the question of whether our experiment captured the entire transgenerational transcriptomic response. First, gene expression has been investigated in only one tissue (i.e., the mantle skirt that synthesises the shell) in this study, whereas in many studies it has been studied in whole organs (*e.g*., Stein et al. 2018) or whole individuals (*e.g*., Hales et al. 2017). This may drastically decrease the number of DEGs detected, capturing only a specific part of the gene expression changes caused by parental environments. However, this tissue-specific approach is particularly relevant in systems such as ours as it allows the specific study of gene expression in the tissue at the origin of phenotypic traits (i.e. shell modifications in this study).

Second, this study investigated only the gene expression in offspring that were not themselves exposed to predator cues, which did not allow us to determine whether the gene expression associated with parental environments is modulated by developmental cues. In *P. acuta*, there is evidence that parental and developmental environments interact to shape the offspring phenotype: the resulting phenotypic patterns showed much stronger TGP in offspring that had not experienced predator cues themselves (WGP masking TGP for most traits; (Luquet & Tariel, 2016)). Although it would be relevant to provide an integrative view of gene expression patterns in all environmental scenarios, our study of DEGs associated with TGP focused on the environmental situation where TGP is the strongest.

Third, the lower number of DEGs in F2 could also result from a time lag between changes in gene expression and antipredator defence induction (Gilbert, 2001). Indeed, the environmental conditions experienced in early life can have important consequences for phenotypes later in life (Gilbert, 2001; Lee et al., 2013) as well as for offspring phenotypes (Burton & Metcalfe, 2014; Tariel-Adam et al., 2023; Yin et al., 2019). For example, maternal provisioning plays a crucial role in transmitting environmental information to offspring by influencing cytoplasmic components, including mRNAs and proteins, during early embryogenesis (Burton & Metcalfe, 2014). This process can impact gene regulation during the maternal-to-zygotic transition, potentially shaping phenotypic responses before they are detectable in adults. In this study, both differential gene expression and antipredator defences in offspring were investigated at the adult stage, and offspring may have integrated the environmental information from their parents early in development engaging them in specific predation risk developmental trajectories. Consequently, the antipredator defence observed in offspring from exposed parents may result from gene expression changes early in development that no longer exist at the adult stage, which could also explain why we did not find a clear common core set of genes between WGP and TGP. For example, Mommer & Bell (2014) showed that maternal experience with predation risk influences genome-wide embryonic expression in *G. aculeatus*, an early expression that may have phenotypic consequences later (Sharda et al., 2021). This complex interplay among the timing of environmental cue perception, gene expression and phenotypic consequences deserves to be explored in more detail, for example by tracking DEGs throughout development.

The functional analysis of DEGs between snails from nonexposed parents and those from exposed parents suggested the same main functions as WGP, i.e., metabolism and transcription regulation. Genes associated with metabolism are mainly involved in the response to oxidative stress and carbohydrate metabolism. One gene might be involved in the regulation of the cytosolic calcium ion concentration, which might influence the shell synthesis as previously mentioned for WGP. Most genes linked with transcription regulation in F2 are involved in methyltransferase activity, suggesting that DNA methylation, a well-known bearer of epigenetic information, may be involved in predator-induced TGP (Fallet et al., 2020). Epigenetic mechanisms, such as DNA methylation and histone modifications, are known to contribute to the transgenerational inheritance of phenotypic traits (Duncan et al., 2014; Fallet et al., 2020 for a review on molluscs). Recent studies on the freshwater gastropod *Potamopyrgus antipodarum* have demonstrated that differential DNA methylation patterns are associated with heritable morphological differentiations among genetically uniform populations (Smithson et al., 2020; Thorson et al., 2017, 2019). These results suggest that the transgenerational inheritance of antipredator traits in *P. acuta* may be mediated by bearers of epigenetic information and call for more investigations of epigenetic inheritance in the context of predator-induced plasticity.

### Comparison of differentially expressed genes associated with WGP and TGP

The number of DEGs associated with TGP was lower than that associated with WGP. As mentioned above, this may have occured because we only investigated DEGs at the adult stage or because F2 snails did not experience direct exposure to predator cues. Surprisingly, the DEGs associated with WGP and TGP did not overlap, suggesting that predator-induced TGP and WGP do not share common genes. This finding is consistent with Hales et al. (2017), who reported that the DEGs for predator-induced WGP and TGP in *D. ambigua* were largely distinct. In contrast, Stein et al. (2018) reported that there were unique sets of genes related to the different forms of plasticity but also identified a common core set of genes in *G. aculeatus*. Despite the absence of common genes in our study, the DEGs in F2 seemed to be involved in the same main functions as those in F1, i.e., metabolism and transcriptional regulation. Consequently, these results suggest that common functions are used for predator-induced WGP and TGP via different genomic pathways. This finding has important implications for the evolution of predator-induced plasticity as it suggests that WGP and TGP can evolve independently.

## Conclusion

This study confirmed predator-induced WGP and TGP in *P. acuta*, showing that antipredator traits (escape behaviour, shell thickness, shell length and shell shape) were induced when snails were exposed during their own development or when they originated from parents exposed to predator cues (but not exposed themselves). These within- and transgenerational phenotypic responses were both associated with differential patterns of gene expression. However, the transcriptomic signal was lower for TGP (a lower number of DEGs), and surprisingly no common genes were shared between WGP and TGP, although the main functions were similar (metabolism and transcription regulation). Expression analysis revealed that only a small set of genes (112 in F1 and 23 in F2) exhibited strong differential expression responses consistently in both generations. Several hypotheses have been proposed to explain these distinct transcriptomic patterns between predator-induced within-generational and transgenerational plasticity. Further research is needed to overcome the limitations of this study by examining transcriptomic signals throughout development, across multiple tissues, and under diverse environmental conditions. This study provides a more integrative view of the underlying molecular mechanisms of WGP and TGP, addressing an important gap in knowledge concerning important processes for adaptation to changing environments.

## Acknowledgements.

We are grateful to Cyril Degletagne who helped in experimental works and to the Rhône-Alpes bioinformatic centre (PRABI) that has supported bioinformatics analyses. This work was financially supported by the BioEnviS research federation (FR3728) of Lyon University and the TEATIME (ANR-21-CE02-0005) grants from the French National Research Agency (ANR).

## Authors Contributions

E.L., S.P. and T.L. conceived the project. J.T., S.P. and E.L. performed the experiments. L.K. performed RNA extractions and libraries. L.D., J.T. and M.H. analysed phenotypic data. L.D., N.S and T.L. performed bioinformatic molecular analyses. L.D. wrote the first version of the manuscript. All authors revised the manuscript.

## Ethics

This work did not require ethical approval from a human subject or animal welfare committee.

## Conflict of interest declaration

We declare we have no competing interests.

## Funding

This work was financially supported by the TEATIME (ANR-21-CE02-0005) grants from the French National Research Agency (ANR).

## Data accessibility

RNA-seq data have been deposited in the European Nucleotide Archive (ENA) under the accession number [ PRJEB87216 ]. The raw sequencing data are publicly available at [ https://www.ebi.ac.uk/ena/browser/view/PRJEB87216 ]. Additionally, scripts used for the analysis, along with the non-genomic data, have been made available on Zenodo. The repository can be accessed at [<u>10.5281/zenodo.15043538</u> ].

## Notes

### Competing Interest Statement

The authors have declared no competing interest.

### Summary of Updates

We have mainly edited english language.

