## Supplementary material for "Transcriptomic basis of within- and trans-generational predator-induced plasticity in the freshwater snail *Physa acuta*"

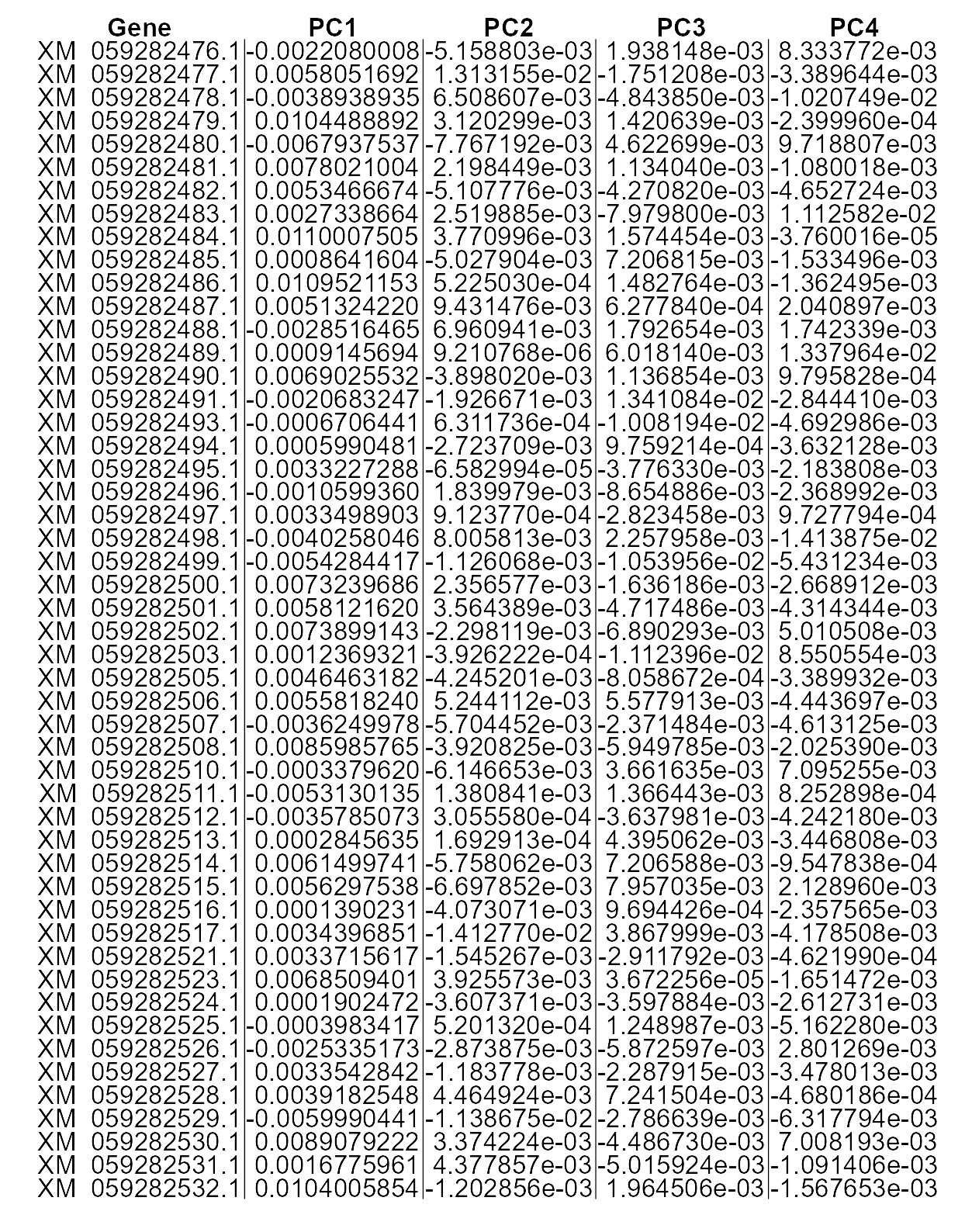

**Figure S1:** Contribution of top genes to the first (PC1) and second (PC2) principal components. Genes with higher loadings have a stronger influence on the variance explained by these components.

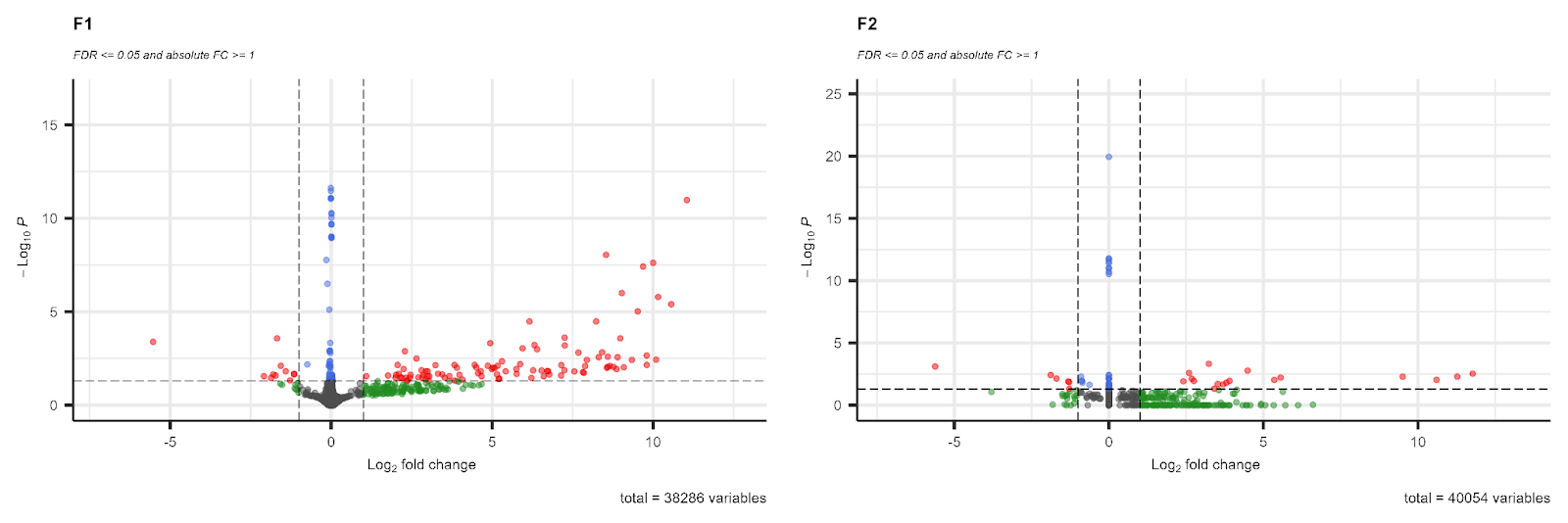

**Figure S2**: Volcano plots showing the distribution of differentially expressed genes between the treatment and control groups for F1 (A) and F2 (B) generations. Log2 Fold Change (log2FC) values are plotted against adjusted p-values (padj). Significance thresholds (padj ≤ 0.05) and fold change criteria (|log2FC| ≥ 1) are indicated by a dotted line (respectively horizontal and vertical); DEGs selected by thresholds are represented red dots.

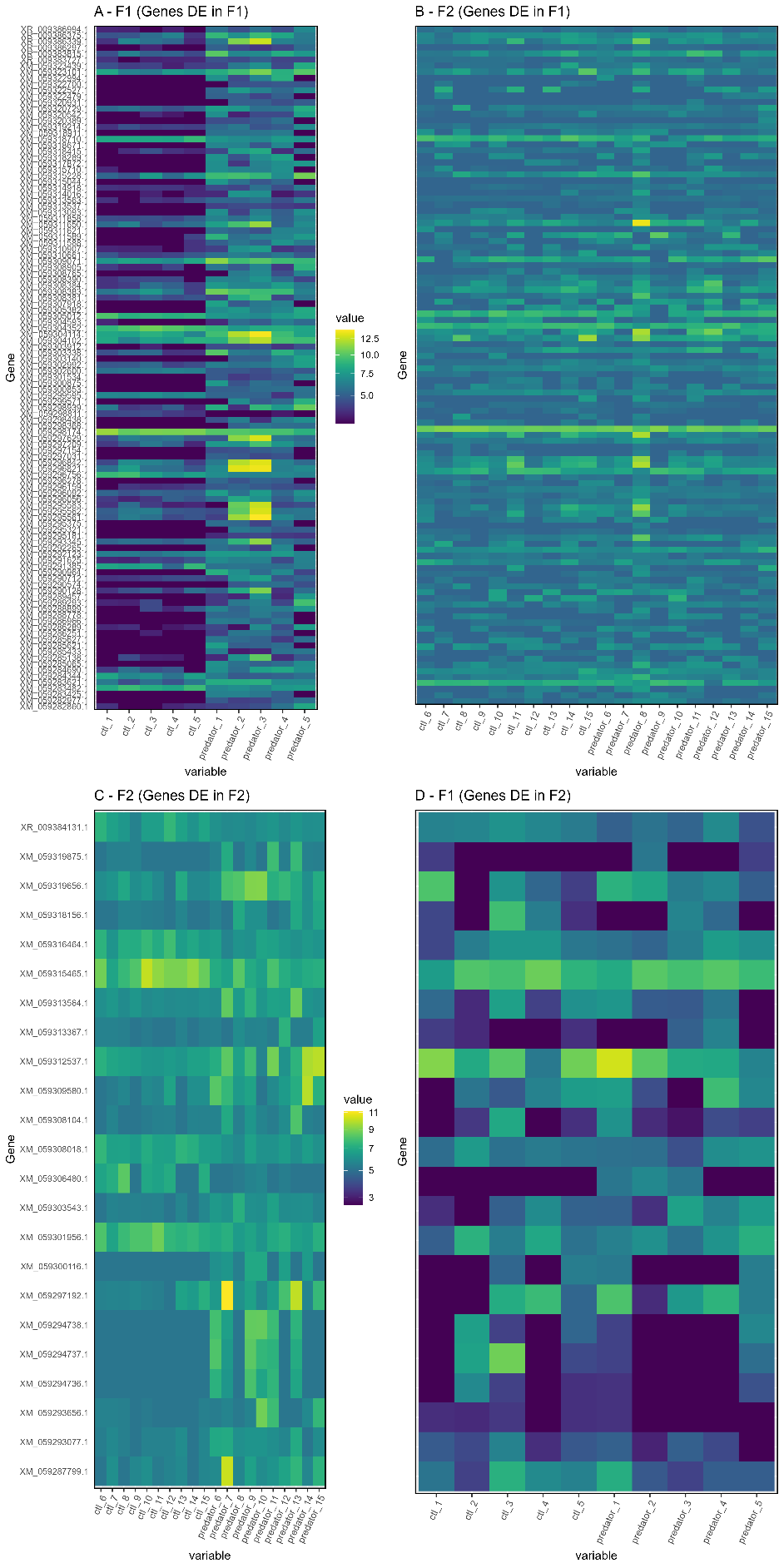

**Figure S3**: Heatmap of read counts of **112** F1 DEGs (top panels) between non-exposed and exposed to predator cues F1 snails (A) and the same genes between F2 snails from non-exposed and exposed parents (B); Heatmap of read counts of **23** F2 DEGs (bottom panels) between non-exposed and exposed to predator cues F2 snails (C) and the same genes between F1 snails from non-exposed and exposed parents (D).

**Table S1:** Detail of the number of snails in each family for each treatment and generation, and survival rates.

| Generation | Family | Treatement | N tot | N | survival (%) |
| --- | --- | --- | --- | --- | --- |
| F1 | 1 | C | 5 | 3 | 60 |
| F1 | 2 | C | 5 | 4 | 80 |
| F1 | 3 | C | 5 | 4 | 80 |
| F1 | 4 | C | 5 | 4 | 80 |
| F1 | 5 | C | 5 | 4 | 80 |
| F1 | 6 | C | 5 | 4 | 80 |
| F1 | 7 | C | 5 | 4 | 80 |
| F1 | 8 | C | 5 | 4 | 80 |
| F1 | 9 | C | 5 | 4 | 80 |
| F1 | 10 | C | 5 | 3 | 60 |
| F1 | 11 | C | 5 | 4 | 80 |
| F1 | 12 | C | 5 | 4 | 80 |
| F1 | 13 | C | 5 | 4 | 80 |
| F1 | 14 | C | 5 | 4 | 80 |
| F1 | 15 | C | 5 | 3 | 60 |
| F1 | 16 | C | 5 | 4 | 80 |
| F1 | 17 | C | 5 | 4 | 80 |
| F1 | 18 | C | 5 | 3 | 60 |
| F1 | 19 | C | 5 | 4 | 80 |
| F1 | 20 | C | 5 | 4 | 80 |
| F1 | 1 | P | 5 | 3 | 60 |
| F1 | 2 | P | 5 | 2 | 40 |
| F1 | 3 | P | 5 | 4 | 80 |
| F1 | 4 | P | 5 | 4 | 80 |
| F1 | 5 | P | 5 | 4 | 80 |
| F1 | 6 | P | 5 | 4 | 80 |
| F1 | 7 | P | 5 | 3 | 60 |
| F1 | 8 | P | 5 | 4 | 80 |
| F1 | 9 | P | 5 | 4 | 80 |
| F1 | 10 | P | 5 | 4 | 80 |
| F1 | 11 | P | 5 | 4 | 80 |
| F1 | 12 | P | 5 | 3 | 60 |
| F1 | 13 | P | 5 | 3 | 60 |
| F1 | 14 | P | 5 | 4 | 80 |
| F1 | 15 | P | 5 | 4 | 80 |
| F1 | 16 | P | 5 | 4 | 80 |
| F1 | 17 | P | 5 | 3 | 60 |
| F1 | 18 | P | 5 | 4 | 80 |
| F1 | 19 | P | 5 | 4 | 80 |
| F1 | 20 | P | 5 | 4 | 80 |
| Generation | Family | Treatement | N tot | N | survival (%) |
| **F2** | 2 | C | 8 | 4 | 50 |
| **F2** | 3 | C | 8 | 5 | 62,5 |
| **F2** | 4 | C | 8 | 5 | 62,5 |
| **F2** | 5 | C | 8 | 4 | 50 |
| **F2** | 6 | C | 8 | 4 | 50 |
| **F2** | 7 | C | 8 | 3 | 37,5 |
| **F2** | 8 | C | 8 | 4 | 50 |
| **F2** | 9 | C | 8 | 3 | 37,5 |
| **F2** | 10 | C | 8 | 4 | 50 |
| **F2** | 11 | C | 8 | 4 | 50 |
| **F2** | 12 | C | 8 | 5 | 62,5 |
| **F2** | 13 | C | 8 | 4 | 50 |
| **F2** | 14 | C | 8 | 5 | 62,5 |
| **F2** | 15 | C | 8 | 6 | 75 |
| **F2** | 16 | C | 8 | 5 | 62,5 |
| **F2** | 17 | C | 8 | 4 | 50 |
| **F2** | 18 | C | 8 | 5 | 62,5 |
| **F2** | 19 | C | 8 | 5 | 62,5 |
| **F2** | 20 | C | 8 | 6 | 75 |
| **F2** | 1 | P | 8 | 6 | 75 |
| **F2** | 2 | P | 8 | 6 | 75 |
| **F2** | 3 | P | 8 | 5 | 62,5 |
| **F2** | 4 | P | 8 | 5 | 62,5 |
| **F2** | 5 | P | 8 | 6 | 75 |
| **F2** | 6 | P | 8 | 6 | 75 |
| **F2** | 7 | P | 8 | 3 | 37,5 |
| **F2** | 8 | P | 8 | 8 | 100 |
| **F2** | 9 | P | 8 | 4 | 50 |
| **F2** | 10 | P | 8 | 4 | 50 |
| **F2** | 11 | P | 8 | 4 | 50 |
| **F2** | 12 | P | 8 | 4 | 50 |
| **F2** | 13 | P | 8 | 3 | 37,5 |
| **F2** | 14 | P | 8 | 5 | 62,5 |
| **F2** | 15 | P | 8 | 6 | 75 |
| **F2** | 16 | P | 8 | 5 | 62,5 |
| **F2** | 17 | P | 8 | 8 | 100 |
| **F2** | 18 | P | 8 | 5 | 62,5 |
| **F2** | 19 | P | 8 | 3 | 37,5 |
| **F2** | 20 | P | 8 | 5 | 62,5 |

**Table S2:** Details of the molecular functions and biological processes of each differentially expressed transcripts in F1 based on the InterPro EMBL-EBI database (Jones et al., 2014; Blum et al., 2024).

| **Gene** | **baseMean** | **log2FoldChange** | **lfcSE** | **pvalue** | **padj** | **Protein family membership** | **biological process** | **molecular function** | **GO terms database** | **Class of function** |
| --- | --- | --- | --- | --- | --- | --- | --- | --- | --- | --- |
| XM_059305382.1 | 72,58 | 11,05 | 3,05 | 2,71524E-15 | 1,05142E-11 | PTHR10694 LYSINE-SPECIFIC DEMETHYLASE | chromatin remodeling (GO:0006338) regulation of gene expression (GO:0010468) regulation of DNA-templated transcription (GO:0006355) | histone demethylase activity (GO:0032452) histone H3K4me/H3K4me2/H3K4me3 demethylase activity (GO:0034647 | PANTHER | Transcription Regulation |
| XM_059290964.1 | 62,54 | 10,56 | 3,26 | 3,96073E-09 | 0 | Mono-ADP-ribosyltransferase ARTD/PARP (IPR052056) | negative regulation of gene expression (GO:0010629) | NAD+ poly-ADP-ribosyltransferase activity (GO:0003950) transcription corepressor activity (GO:0003714) | InterPro/PANTHER | Transcription Regulation |
| XM_059317872.1 | 47,2 | 10,15 | 3,14 | 1,55403E-09 | 0 | GTPase GIMA/IAN/Toc (IPR045058) |  | GTP binding (GO:0005525) | InterPro | Metabolism |
| XM_059322994.1 | 62,49 | 10,09 | 3,67 | 7,29422E-06 | 0,00366 |  |  |  |  | Unknown |
| XM_059322527.1 | 39,69 | 10 | 3,01 | 1,96594E-11 | 2,42222E-08 | Dynein heavy chain (IPR026983) | microtubule-based movement (GO:0007018) | ATP binding (GO:0005524) dynein light intermediate chain binding (GO:0051959) dynein intermediate chain binding (GO:0045505) minus-end-directed microtubule motor activity (GO:0008569) ATP hydrolysis activity (GO:0016887) | InterPro | Microtubule Motor activity |
| XM_059299571.1 | 55,28 | 9,8 | 3,71 | 1,71962E-05 | 0,007 |  |  |  |  | Unknown |
| XM_059318289.1 | 48,89 | 9,8 | 3,51 | 3,95201E-06 | 0,00219 | Rho-associated Serine/Threonine Kinase (IPR050839) | protein phosphorylation (GO:0006468) | ATP binding (GO:0005524) protein kinase activity (GO:0004672) protein serine/threonine kinase activity (GO:0004674) | InterPro | Signal transduction |
| XM_059285065.1 | 33,25 | 9,69 | 2,96 | 3,23349E-11 | 3,81074E-08 | PTHR46232 SMARCE1 REGULATOR OF CHROMATIN | negative regulation of DNA-templated transcription (GO:0045892) | nuclear receptor binding (GO:0016922) nucleosomal DNA binding (GO:0031492) | InterPro/PANTHER | Transcription regulation |
| XM_059313093.1 | 32,69 | 9,52 | 3,08 | 1,03156E-08 | 0,00001 | Serpin family (IPR000215) | serine-type endopeptidase inhibitor activity (GO:0004867) |  | InterPro | Immune/Defense response |
| XM_059307918.1 | 37,57 | 9,34 | 3,46 | 7,75519E-06 | 0,00378 | Regulatory associated protein of TOR (IPR004083) | regulation of autophagy (GO:0010506) cellular response to starvation (GO:0009267) TORC1 signaling (GO:0038202) cellular response to amino acid stimulus (GO:0071230) positive regulation of cell growth (GO:0030307) TOR signaling (GO:0031929) | protein-macromolecule adaptor activity (GO:0030674) protein binding (GO:0005515) | InterPro/PANTHER | Cell growth regulation |
| XM_059303140.1 | 35,23 | 9,09 | 3,57 | 2,77026E-05 | 0,00939 | STE20/SPS1-related Proline-Alanine-rich Kinase (IPR050629) Mitogen-activated protein (MAP) kinase kinase kinase kinase (IPR021160) | intracellular signal transduction (GO:0035556) protein phosphorylation (GO:0006468) | protein serine/threonine kinase activity (GO:0004674) MAP kinase kinase kinase kinase activity (GO:0008349) | PANTHER | Signal transduction |
| XM_059311621.1 | 22,77 | 9,03 | 2,91 | 9,21574E-10 | 9,99207E-07 | Protein O-mannosyl-transferase TMTC (IPR052346) | endoplasmic reticulum unfolded protein response (GO:0030968) protein O-linked mannosylation (GO:0035269) | mannosyltransferase activity (GO:0000030) protein binding (GO:0005515) | InterPro/PANTHER | Intracellular transport |
| XM_059298438.1 | 25,18 | 8,98 | 3,12 | 3,31012E-07 | 0,00026 |  |  | metal ion binding (GO:0046872) | InterPro | Ion binding/transport |
| XM_059318671.1 | 26,76 | 8,89 | 3,3 | 5,14503E-06 | 0,0027 | Proteasome, subunit alpha/beta (IPR001353) Proteasome alpha-type subunit (IPR023332) | ubiquitin-dependent protein catabolic process (GO:0006511) proteolysis involved in protein catabolic process (GO:0051603) |  | InterPro | Metabolism |
| XM_059289457.1 | 30,98 | 8,86 | 3,55 | 3,80749E-05 | 0,01158 |  |  |  |  | Unknown |
| XM_059285621.1 | 27,8 | 8,77 | 3,46 | 2,59408E-05 | 0,0089 | UNC-93-like transmembrane regulator (IPR051617) Ion channel regulatory protein, UNC-93 (IPR010291) |  |  | InterPro | Transmembrane transport |
| XM_059315044.1 | 25,41 | 8,66 | 3,4 | 2,18185E-05 | 0,00789 | PTHR24409 C2H2-type zinc-finger domain-containing protein | regulation of transcription by RNA polymerase II (GO:0006357) | RNA polymerase II transcription regulatory region sequence-specific DNA binding (GO:0000977) DNA-binding transcription factor activity, RNA polymerase II-specific (GO:0000981) | InterPro | Transcription regulation |
| XM_059283425.1 | 21,66 | 8,59 | 3,21 | 4,70019E-06 | 0,00255 | PTHR24347 Calcium/calmodulin-dependent protein kinase | protein phosphorylation (GO:0006468) | protein kinase activity (GO:0004672) ATP binding (GO:0005524) | InterPro | Signal transduction |
| XM_059308765.1 | 24,49 | 8,59 | 3,4 | 2,56841E-05 | 0,0089 | Galectin-like (IPR044156) |  | carbohydrate binding (GO:0030246) | InterPro | Extracellular matrix organisation |
| XM_059300875.1 | 24,55 | 8,56 | 3,42 | 3,11826E-05 | 0,00994 | ATP-binding cassette transporter C-like (IPR050173) | transmembrane transport (GO:0055085) | ATP hydrolysis activity (GO:0016887) ATP binding (GO:0005524) ATPase-coupled transmembrane transporter activity (GO:0042626) ABC-type transporter activity (GO:0140359) | InterPro | Transmembrane transport |
| XM_059300853.1 | 15,75 | 8,54 | 2,72 | 6,66461E-12 | 9,03254E-09 | Sulfhydryl oxidase ALR/ERV (IPR039799) |  | thiol oxidase activity (GO:0016972) flavin-dependent sulfhydryl oxidase activity (GO:0016971) | InterPro | Metabolism |
| XM_059318911.1 | 18,53 | 8,42 | 3,11 | 2,46142E-06 | 0,00145 | Ras-like guanine nucleotide exchange factor (IPR008937) | small GTPase-mediated signal transduction (GO:0007264) signal transduction (GO:0007165) | guanyl-nucleotide exchange factor activity (GO:0005085) protein binding (GO:0005515) | InterPro | Signal transduction |
| XM_059286966.1 | 17,97 | 8,31 | 3,14 | 5,18737E-06 | 0,0027 | Spindle and centriole-associated protein 1 (IPR031387) | metaphase chromosome alignment (GO:0051310) mitotic spindle assembly (GO:0090307) regulation of centriole replication (GO:0046599) |  | InterPro/PANTHER | Cell division regulation |
| XM_059285627.1 | 14,63 | 8,23 | 2,85 | 3,69146E-08 | 0,00003 |  | regulation of transcription by RNA polymerase II (GO:0006357) | RNA polymerase II cis-regulatory region sequence-specific DNA binding (GO:0000978) DNA-binding transcription factor activity, RNA polymerase II-specific (GO:0000981) transcription cis-regulatory region binding (GO:0000976) | InterPro/PANTHER | Transcription regulation |
| XM_059288778.1 | 14,3 | 7,94 | 3,08 | 7,80906E-06 | 0,00378 | Peptidase C14 family (IPR002398) | proteolysis (GO:0006508) | cysteine-type endopeptidase activity (GO:0004197) cysteine-type peptidase activity (GO:0008234) | InterPro | Immune/Defense response |
| XM_059298368.1 | 14,94 | 7,88 | 3,17 | 2,25502E-05 | 0,00804 | Zinc finger (IPR050331) | regulation of gene expression (GO:0010468) |  | InterPro | Transcription regulation |
| XM_059292265.1 | 16,12 | 7,85 | 3,33 | 7,18635E-05 | 0,01855 |  | protein modification by small protein removal (GO:0070646) protein deubiquitination (GO:0016579) | deubiquitinase activity (GO:0101005) | PANTHER | Metabolism |
| XM_059285433.1 | 15,92 | 7,82 | 3,31 | 6,4531E-05 | 0,01682 | Tyrosine-protein kinase BAZ1B (IPR047174) | DNA damage response (GO:0006974) chromatin remodeling (GO:0006338) | histone kinase activity (GO:0035173) histone binding (GO:0042393) | PANTHER | Transcription Regulation |
| XM_059301534.1 | 12,07 | 7,68 | 2,93 | 2,65355E-06 | 0,00153 | Complex I intermediate-associated protein 30, mitochondrial (IPR039131) | mitochondrial respiratory chain complex I assembly (GO:0032981) NADH dehydrogenase complex assembly (GO:0010257) mitochondrial electron transport, NADH to ubiquinone (GO:0006120) | unfolded protein binding (GO:0051082) | PANTHER | Metabolism |
| XM_059295321.1 | 12,82 | 7,55 | 3,2 | 5,87782E-05 | 0,01557 |  |  |  |  | Unknown |
| XM_059282977.1 | 8,9 | 7,26 | 2,76 | 9,08946E-07 | 0,00063 | Septin (IPR016491) |  | GTP binding (GO:0005525) | InterPro | Signal transduction |
| XM_059315710.1 | 38,49 | 7,25 | 1,87 | 2,86204E-07 | 0,00024 | Ubiquitin carboxyl-terminal hydrolase (IPR050185) | protein deubiquitination (GO:0016579) | zinc ion binding (GO:0008270) cysteine-type deubiquitinase activity (GO:0004843) | InterPro | Metabolism |
| XM_059295375.1 | 10,14 | 7,24 | 3,07 | 4,65992E-05 | 0,01358 | SCF Complex F-box/Wd Repeat-Containing Protein (IPR052301) | SCF-dependent proteasomal ubiquitin-dependent protein catabolic process (GO:0031146) | protein binding (GO:0005515) | InterPro/PANTHER | Proteolysis/Protein degradation |
| XM_059296278.1 | 8,83 | 7,15 | 2,93 | 1,78811E-05 | 0,00713 | Zinc finger CCHC-type and RNA-binding motif-containing protein 1 (IPR044598) | mRNA splicing, via spliceosome (GO:0000398) | zinc ion binding (GO:0008270) nucleic acid binding (GO:0003676) RNA binding (GO:0003723) | InterPro | Transcription Regulation |
| XM_059290574.1 | 10,32 | 7,14 | 3,18 | 0,000118375 | 0,02567 | Zinc finger ZZ-type and EF-hand domain-containing protein 1 (IPR040099) |  | zinc ion binding (GO:0008270) | InterPro | Metabolism |
| XM_059320389.1 | 7,77 | 6,77 | 3,02 | 9,77958E-05 | 0,02285 | Piezo family (IPR027272) |  | mechanosensitive monoatomic ion channel activity (GO:0008381) | InterPro | Signal transduction |
| XM_059322479.1 | 7,26 | 6,74 | 2,93 | 5,39323E-05 | 0,01523 | Mitochondrial glycine transporter Hem25/SLC25A38 (IPR030847) Mitochondrial carrier protein (IPR002067) | glycine import into mitochondrion (GO:1904983) transmembrane transport (GO:0055085) | glycine transmembrane transporter activity (GO:0015187) | InterPro | Metabolism |
| XM_059297154.1 | 6,98 | 6,71 | 2,91 | 5,23144E-05 | 0,01493 | Transmembrane protein 272 (IPR040350) |  |  |  | Transmembrane transport |
| XM_059320931.1 | 7,26 | 6,6 | 3,02 | 0,000133354 | 0,02779 | PTHR43201 ATP-dependent AMP-binding enzyme | fatty acid metabolic process (GO:0006631) | medium-chain fatty acid-CoA ligase activity (GO:0031956) | InterPro/PANTHER | Metabolism |
| XM_059297031.1 | 6,28 | 6,54 | 2,85 | 4,7763E-05 | 0,01377 | Cytochrome P450, E-class, group I (IPR002401) |  | heme binding (GO:0020037) iron ion binding (GO:0005506) monooxygenase activity (GO:0004497) oxidoreductase activity, acting on paired donors, with incorporation or reduction of molecular oxygen (GO:0016705) | InterPro | Metabolism |
| XM_059295561.1 | 1903,92 | 6,39 | 1,59 | 1,53872E-06 | 0,00102 |  |  |  |  | Unknown |
| XM_059311589.1 | 83,03 | 6,32 | 1,54 | 8,45596E-07 | 0,0006 | IPR001752 Kinesin motor domain | microtubule-based movement (GO:0007018) | ATP binding (GO:0005524) microtubule motor activity (GO:0003777) microtubule binding (GO:0008017) protein binding (GO:0005515) | InterPro | Cell division regulation |
| XM_059295583.1 | 771 | 6,27 | 2,12 | 4,63E-05 | 0,01358 |  |  |  |  | Unknown |
| XM_059322700.1 | 5,82 | 6,22 | 2,95 | 0,000190373 | 0,03487 | Neurotransmitter-gated ion-channel (IPR006201) | monoatomic ion transport (GO:0006811) monoatomic ion transmembrane transport (GO:0034220) | extracellular ligand-gated monoatomic ion channel activity (GO:0005230) monoatomic ion channel activity (GO:0005216) transmembrane signaling receptor activity (GO:0004888) | InterPro | Signal transduction |
| XM_059311588.1 | 29,32 | 6,16 | 1,35 | 3,75726E-08 | 0,00003 |  | microtubule-based movement (GO:0007018) | protein binding (GO:0005515) ATP binding (GO:0005524) microtubule motor activity (GO:0003777) microtubule binding (GO:0008017) | InterPro | Cell division regulation |
| XM_059304593.1 | 14,97 | 5,94 | 1,68 | 1,32973E-06 | 0,0009 | Sodium leak channel NALCN (IPR028823) | monoatomic ion transport (GO:0006811) transmembrane transport (GO:0055085) | monoatomic ion channel activity (GO:0005216) monoatomic cation channel activity (GO:0005261) | InterPro | Ion transport |
| XM_059313537.1 | 3,62 | 5,87 | 2,53 | 1,43217E-05 | 0,00636 | Ran binding protein RanBP1-like (IPR045255) | intracellular transport (GO:0046907) |  | InterPro | Transport |
| XM_059285138.1 | 139,7 | 5,76 | 2,04 | 8,50802E-05 | 0,02097 | Dermatopontin (IPR026645) | collagen fibril organization (GO:0030199) |  | InterPro | Extracellular matrix organization |
| XM_059320542.1 | 35,15 | 5,75 | 1,77 | 3,75528E-05 | 0,01157 | Serine/Threonine Dehydratase (IPR050147) | isoleucine biosynthetic process (GO:0009097) L-serine catabolic process (GO:0006565) threonine catabolic process (GO:0006567) | threonine deaminase activity (GO:0004794) L-serine ammonia-lyase activity (GO:0003941) | PANTHER | Metabolism |
| XM_059286251.1 | 12,32 | 5,42 | 1,93 | 5,6433E-05 | 0,01534 |  | proteasome-mediated ubiquitin-dependent protein catabolic process (GO:0043161) | ubiquitin binding (GO:0043130) | PANTHER | Proteolysis/Protein degradation |
| XM_059296821.1 | 2195,19 | 5,3 | 1,49 | 9,70307E-06 | 0,00446 |  |  |  |  | Unknown |
| XM_059314016.1 | 26,77 | 5,22 | 2,16 | 0,000205243 | 0,03734 | Annexin (IPR001464) |  | calcium-dependent phospholipid binding (GO:0005544) calcium ion binding (GO:0005509) | InterPro | Ion transport |
| XM_059295181.1 | 2,78 | 5,2 | 2,59 | 0,000222315 | 0,03965 | PTHR24379 C2H2-type zinc-finger domain-containing protein | regulation of transcription by RNA polymerase II (GO:0006357) | RNA polymerase II transcription regulatory region sequence-specific DNA binding (GO:0000977) DNA-binding transcription factor activity, RNA polymerase II-specific (GO:0000981) | InterPro | Gene expression regulation |
| XM_059297629.1 | 1121,03 | 5,17 | 1,55 | 2,04049E-05 | 0,00778 |  |  |  |  | Unknown |
| XM_059289293.1 | 88,33 | 5,15 | 1,82 | 9,22596E-05 | 0,02231 | PTHR36496 CADHERIN DOMAIN-CONTAINING PROTEIN |  |  | PANTHER | Cell adhesion |
| XM_059296842.1 | 1447,78 | 5,08 | 1,59 | 3,24117E-05 | 0,01022 |  |  |  |  | Unknown |
| XM_059295562.1 | 964,7 | 5,02 | 1,57 | 3,05138E-05 | 0,00985 | PTHR36912 ANTIGEN 332, DBL-LIKE PROTEIN-RELATED |  |  | PANTHER | Unknown |
| XR_009386339.1 | 908,13 | 4,99 | 1,59 | 3,84419E-05 | 0,01158 |  |  |  |  | Unknown |
| XR_009386994.1 | 19,76 | 4,94 | 1,1 | 6,55838E-07 | 0,00048 |  |  |  |  | Unknown |
| XR_009386297.1 | 7,94 | 4,86 | 1,64 | 2,09444E-05 | 0,00778 |  |  |  |  | Unknown |
| XM_059286289.1 | 33,47 | 4,67 | 1,7 | 0,000137617 | 0,02798 | Fibrinogen C-terminal domain-containing protein (IPR050373) |  |  | InterPro | Extracellular matrix organization |
| XM_059310907.1 | 33,61 | 4,65 | 1,53 | 5,715E-05 | 0,01534 | PTHR46147 Histone-lysine N-methyltransferase SET2 subfamily | regulation of DNA-templated transcription (GO:0006355) | DNA binding (GO:0003677) protein binding (GO:0005515) chromatin binding (GO:0003682) histone methyltransferase activity (GO:0042054) | InterPro | Gene expression regulation |
| XM_059282860.1 | 37,28 | 4,57 | 1,55 | 7,64667E-05 | 0,01955 | Peptidase C2, calpain family (IPR022684) | proteolysis (GO:0006508) | calcium-dependent cysteine-type endopeptidase activity (GO:0004198) | InterPro | Proteolysis/Protein degradation |
| XM_059308905.1 | 59,71 | 4,51 | 1,35 | 2,8406E-05 | 0,00951 | Macin (IPR029230) | defense response (GO:0006952) |  | InterPro | Immune/Defense response |
| XM_059303912.1 | 5,24 | 4,46 | 1,47 | 1,70744E-05 | 0,007 |  |  |  |  | Unknown |
| XM_059290128.1 | 246,29 | 4,07 | 1,64 | 0,000242469 | 0,0424 | Multicopper oxidase (IPR045087) | iron ion transport (GO:0006826) obsolete iron ion homeostasis (GO:0055072) | copper ion binding (GO:0005507) oxidoreductase activity (GO:0016491) | InterPro/PANTHER | Ion transport |
| XM_059298939.1 | 251,4 | 3,98 | 1,4 | 0,000103623 | 0,02367 | Fibrinogen C-terminal domain-containing protein (IPR050373) |  |  | InterPro | Extracellular matrix organization |
| XM_059311650.1 | 427,06 | 3,9 | 1,2 | 3,04627E-05 | 0,00985 |  |  |  |  | Unknown |
| XM_059318415.1 | 59,31 | 3,83 | 1,55 | 0,000286974 | 0,04743 | PTHR10127 Astacin-like Metalloprotease | proteolysis (GO:0006508) | metallopeptidase activity (GO:0008237) zinc ion binding (GO:0008270) metalloendopeptidase activity (GO:0004222) | InterPro/PANTHER | Proteolysis/Protein degradation |
| XM_059303338.1 | 173,18 | 3,83 | 1,11 | 1,69775E-05 | 0,007 |  |  |  |  | Unknown |
| XM_059293345.1 | 337,86 | 3,67 | 1,33 | 0,000134619 | 0,02779 | PTHR24039 Extracellular Matrix and Cell Adhesion Regulators |  | protein binding (GO:0005515) calcium ion binding (GO:0005509) | InterPro/PANTHER | Cell adhesion |
| XM_059308381.1 | 119,55 | 3,52 | 1,32 | 0,000179123 | 0,03348 | Extracellular Matrix Assembly and Organization (IPR050525) |  |  | InterPro | Extracellular matrix organization |
| XM_059323439.1 | 55,07 | 3,45 | 1,19 | 9,29963E-05 | 0,02231 | PTHR43884 Acyl-CoA dehydrogenase-related enzymes |  | acyl-CoA dehydrogenase activity (GO:0003995) oxidoreductase activity, acting on the CH-CH group of donors (GO:0016627) flavin adenine dinucleotide binding (GO:0050660) | InterPro/PANTHER | Metabolism |
| XM_059304114.1 | 2058,42 | 3,32 | 1,12 | 8,33356E-05 | 0,02072 | Aerolysin-like pore-forming protein (IPR053280) |  |  | InterPro | Unknown |
| XR_009383815.1 | 190,21 | 3,24 | 0,94 | 1,83908E-05 | 0,00722 |  |  |  |  | Unknown |
| XM_059297209.1 | 136,91 | 3,05 | 1,08 | 0,000130886 | 0,02779 |  |  | chitin binding (GO:0008061) | InterPro | Carbohydrate binding |
| XM_059288899.1 | 42,57 | 3,04 | 1,24 | 0,000313671 | 0,04972 | NAD(P)-dependent epimerase/dehydratase (IPR051225) | threonine catabolic process (GO:0006567) | L-threonine 3-dehydrogenase activity (GO:0008743) | InterPro | Metabolism |
| XM_059296056.1 | 62,31 | 3 | 1,05 | 0,000135324 | 0,02779 | GTPase GIMA/IAN/Toc (IPR045058) |  | GTP binding (GO:0005525) | InterPro | Signal transduction |
| XR_009386375.1 | 172,69 | 3 | 0,96 | 5,48765E-05 | 0,01533 |  |  |  |  | Unknown |
| XM_059290712.1 | 46,12 | 2,94 | 1,12 | 0,000244596 | 0,04247 | Calmodulin/Myosin light chain/Troponin C-like (IPR050230) |  | calcium ion binding (GO:0005509) | InterPro | Calcium signaling |
| XM_059308390.1 | 36,67 | 2,94 | 0,92 | 5,69491E-05 | 0,01534 |  |  |  |  | Unknown |
| XM_059308383.1 | 405,39 | 2,86 | 0,99 | 0,000110956 | 0,02445 | Extracellular Matrix Assembly and Organization (IPR050525) |  |  | InterPro | Extracellular matrix organization |
| XM_059284690.1 | 84,62 | 2,84 | 1,03 | 0,000165399 | 0,03225 |  |  |  |  | Unknown |
| XM_059291625.1 | 35,95 | 2,8 | 0,88 | 4,52666E-05 | 0,01348 | GTPase GIMA/IAN/Toc (IPR045058) Hedgehog protein (IPR001657) | multicellular organism development (GO:0007275) cell-cell signaling (GO:0007267) protein autoprocessing (GO:0016540) | GTP binding (GO:0005525) | InterPro | Signal transduction |
| XM_059315228.1 | 413,99 | 2,71 | 0,99 | 0,000168739 | 0,03244 |  |  |  |  | Unknown |
| XM_059283621.1 | 148,93 | 2,65 | 0,71 | 6,16735E-06 | 0,00315 | PTHR45632 Kelch-like and N-acetylneuraminate epimerase |  | protein binding (GO:0005515) | PANTHER | Protein binding |
| XM_059308384.1 | 67,34 | 2,46 | 0,84 | 0,000105924 | 0,0237 | Extracellular Matrix Assembly and Organization (IPR050525) |  |  | InterPro | Extracellular matrix organization |
| XM_059304102.1 | 1004,42 | 2,44 | 0,88 | 0,000174762 | 0,03336 | Transglutaminase-like Superfamily Modulators (IPR053041) |  |  | InterPro | Protein modification |
| XM_059314918.1 | 14,43 | 2,37 | 0,9 | 0,000282736 | 0,04706 | PTHR24412 Kelch-like E3 ubiquitin ligase complex adapters |  | protein binding (GO:0005515) | PANTHER | Protein binding |
| XR_009383727.1 | 6,5 | 2,36 | 0,85 | 0,000298873 | 0,04819 |  |  |  |  | Unknown |
| XM_059313563.1 | 17,94 | 2,31 | 0,85 | 0,000182239 | 0,03383 |  |  | protein binding (GO:0005515) | InterPro | Protein binding |
| XM_059309071.1 | 643,3 | 2,29 | 0,57 | 2,15269E-06 | 0,0013 | Extracellular Matrix Assembly and Organization (IPR050525) |  |  | InterPro | Extracellular matrix organization |
| XM_059323101.1 | 505,89 | 2,25 | 0,69 | 3,68924E-05 | 0,01149 | Proton-linked Monocarboxylate Transporter (IPR050327) Major facilitator superfamily (IPR011701) | transmembrane transport (GO:0055085) | transmembrane transporter activity (GO:0022857) | InterPro | Transport |
| XM_059319214.1 | 25,3 | 2,19 | 0,8 | 0,000176552 | 0,03347 | Fibrinogen C-terminal domain-containing protein (IPR050373) |  |  | InterPro | Extracellular matrix organization |
| XM_059320729.1 | 91,8 | 2,14 | 0,78 | 0,000164567 | 0,03225 |  |  |  |  | Unknown |
| XM_059302992.1 | 53,73 | 2,1 | 0,7 | 8,27754E-05 | 0,02072 | Mono-ADP-ribosyltransferase and antiviral protein (IPR051712) | obsolete protein mono-ADP-ribosylation (GO:0140289) | NAD+ poly-ADP-ribosyltransferase activity (GO:0003950) NAD+-protein ADP-ribosyltransferase activity (GO:1990404) | InterPro/PANTHER | Metabolism |
| XM_059311858.1 | 37,18 | 2,06 | 0,59 | 1,7301E-05 | 0,007 | Auxin-regulated embryogenesis mediator (IPR052957) |  |  | InterPro | Development |
| XM_059296092.1 | 69,92 | 2,01 | 0,68 | 0,000104807 | 0,02367 | GTPase GIMA/IAN/Toc (IPR045058) |  | GTP binding (GO:0005525) | InterPro | Signal transduction |
| XM_059296159.1 | 9,47 | 2 | 0,69 | 0,000178966 | 0,03348 |  |  |  |  | Unknown |
| XM_059310661.1 | 25,15 | 1,77 | 0,6 | 0,000117771 | 0,02567 |  |  |  |  | Unknown |
| XM_059292123.1 | 106,71 | 1,09 | 0,37 | 0,000133868 | 0,02779 | Fibrinogen C-terminal domain-containing protein (IPR050373) |  |  | InterPro | Extracellular matrix organization |
| XM_059298174.1 | 1392,25 | -1,15 | 0,38 | 9,58988E-05 | 0,0226 | GRB10-interacting GYF (IPR051640) |  | protein binding (GO:0005515) | InterPro | Protein binding |
| XM_059318710.1 | 231,36 | -1,15 | 0,37 | 8,09389E-05 | 0,0205 | Type 1 protein exporter (IPR039421) | transmembrane transport (GO:0055085) | ATP binding (GO:0005524) ATP hydrolysis activity (GO:0016887) ABC-type transporter activity (GO:0140359) | InterpPro | Transport |
| XM_059304352.1 | 624,91 | -1,27 | 0,48 | 0,000283 | 0,04706 | DNA mismatch repair protein MutL/Mlh/PMS (IPR002099) | mismatch repair (GO:0006298) | ATP binding (GO:0005524) mismatched DNA binding (GO:0030983) ATP hydrolysis activity (GO:0016887) ATP-dependent DNA damage sensor activity (GO:0140664) | InterpPro | DNA repair |
| XM_059305012.1 | 343,89 | -1,41 | 0,45 | 5,63982E-05 | 0,01534 | Store-operated calcium entry-associated regulatory factor (IPR009567) | regulation of store-operated calcium entry (GO:2001256) |  | InterpPro | Calcium signaling |
| XM_059283482.1 | 278,67 | -1,57 | 0,45 | 2,09007E-05 | 0,00778 |  |  | protein binding (GO:0005515) | InterPro | Protein binding |
| XM_059302600.1 | 37,66 | -1,68 | 0,38 | 3,2529E-07 | 0,00026 | Short-chain dehydrogenase/reductase SDR (IPR002347) |  | oxidoreductase activity (GO:0016491) | InterPro | Metabolism |
| XM_059299595.1 | 59,13 | -1,72 | 0,6 | 0,000119856 | 0,02578 |  |  |  |  | Unknown |
| XM_059296756.1 | 225,89 | -1,8 | 0,6 | 9,51935E-05 | 0,0226 | Small GTPase (IPR001806) |  | GTP binding (GO:0005525) GTPase activity (GO:0003924) | InterPro | Signal transduction |
| XM_059284344.1 | 89,96 | -1,87 | 0,7 | 0,000189759 | 0,03487 |  |  |  |  | Unknown |
| XM_059291385.1 | 153,86 | -2,09 | 0,74 | 0,000131749 | 0,02779 | DNA damage-induced apoptosis suppressor protein (IPR043522) |  | negative regulation of intrinsic apoptotic signaling pathway in response to DNA damage (GO:1902230) | InterPro | Apoptosis regulation |
| XM_059298811.1 | 3,83 | -5,53 | 1,5 | 5,2448E-07 | 0,00041 |  |  |  |  | Unknown |

**Table S3:** Details of the molecular functions and biological processes of each differentially expressed transcripts in F2 based on the InterPro EMBL-EMI database (Jones et al., 2014; Blum et al., 2024).

| **Gene** | **baseMean** | **l2FC** | **lfcSE** | | **pvalue** | **padj** | **Protein family membership** | **biological process** | **molecular function** | **GO terms database** | **Class of function** |
| --- | --- | --- | --- | --- | --- | --- | --- | --- | --- | --- | --- |
| XM_059294738.1 | 53,09 | 11,77 | 3,91 | 1,8884E-06 | | 0,0029 | Cytosine-specific C5-methyltransferase (IPR050750) |  | methyltransferase activity (GO:0008168) | InterPro | Transcription regulation |
| XM_059294737.1 | 39,45 | 11,27 | 3,87 | 4,38371E-06 | | 0,005 | Cytosine-specific C5-methyltransferase (IPR050750) |  | methyltransferase activity (GO:0008168) | InterPro | Transcription regulation |
| XM_059294736.1 | 26,39 | 10,6 | 3,8 | 1,17623E-05 | | 0,0091 | Cytosine-specific C5-methyltransferase (IPR050750) |  | methyltransferase activity (GO:0008168) | InterPro | Transcription regulation |
| XM_059300116.1 | 11,45 | 9,51 | 3,4 | 4,72867E-06 | | 0,0051 | C2H2-type Zinc-Finger Transcription Regulators (IPR050717) | embryo development (GO:0009790) regulation of transcription by RNA polymerase II (GO:0006357 | RNA polymerase II transcription regulatory region sequence-specific DNA binding (GO:0000977) DNA-binding transcription factor activity, RNA polymerase II-specific (GO:0000981) | PANTHER | Transcription regulation |
| XM_059293656.1 | 33 | 5,56 | 1,49 | 5,85808E-06 | | 0,006 |  |  | protein binding (GO:0005515) DNA binding (GO:0003677) | InterPro | Protein binding |
| XM_059319875.1 | 19,08 | 5,35 | 1,73 | 1,23407E-05 | | 0,0092 |  | carbohydrate derivative metabolic process (GO:1901135) | carbohydrate derivative binding (GO:0097367) | InterPro | Metabolism |
| XM_059287799.1 | 75,51 | 4,49 | 1,07 | 9,28029E-07 | | 0,0017 | Respiratory burst oxidase/Ferric reductase (IPR050369) Haem peroxidase, animal-type (IPR019791) | response to oxidative stress (GO:0006979) superoxide anion generation (GO:0042554) defense response (GO:0006952) | peroxidase activity (GO:0004601) heme binding (GO:0020037) oxidoreductase activity (GO:0016491) calcium ion binding (GO:0005509) | InterPro/PANTHER | Metabolism |
| XM_059318156.1 | 11,99 | 3,9 | 1,23 | 1,66176E-05 | | 0,0111 | Glucose-methanol-choline oxidoreductase (IPR012132) |  | oxidoreductase activity, acting on CH-OH group of donors (GO:0016614) flavin adenine dinucleotide binding (GO:0050660) | InterPro | Metabolism |
| XM_059308104.1 | 16,72 | 3,8 | 1,13 | 2,72889E-05 | | 0,0155 |  |  |  |  | Unknown |
| XM_059297192.1 | 189,03 | 3,69 | 1,17 | 4,13486E-05 | | 0,0213 |  |  | chitin binding (GO:0008061) | InterPro | Chitin binding |
| XM_059309580.1 | 82,82 | 3,52 | 1,1 | 3,78143E-05 | | 0,0199 |  |  | heme binding (GO:0020037) oxygen binding (GO:0019825) | InterPro | Oxygen binding |
| XM_059313367.1 | 8,73 | 3,41 | 1,22 | 0,000120042 | | 0,048 | Amidophosphoribosyltransferase (IPR005854) | purine nucleobase biosynthetic process (GO:0009113) | amidophosphoribosyltransferase activity (GO:0004044) | InterPro | Purine biosynthesis |
| XM_059303543.1 | 12,84 | 3,23 | 0,71 | 2,18495E-07 | | 0,0005 |  |  |  |  | Unknown |
| XM_059293077.1 | 15,45 | 2,75 | 0,8 | 1,69131E-05 | | 0,0111 |  |  |  |  | Unknown |
| XM_059319656.1 | 96,55 | 2,68 | 0,74 | 9,21567E-06 | | 0,0075 | Interferon alpha-inducible protein IFI6/IFI27-like (IPR009311) |  |  | InterPro | Immune response |
| XM_059313564.1 | 51,45 | 2,59 | 0,64 | 1,52502E-06 | | 0,0025 | Peptidase M1, alanine aminopeptidase/leukotriene A4 hydrolase (IPR001930) Aminopeptidase N-type (IPR034016) | proteolysis (GO:0006508) | metallopeptidase activity (GO:0008237) zinc ion binding (GO:0008270) | InterPro | Metabolism |
| XM_059312537.1 | 158,18 | 2,41 | 0,7 | 1,89922E-05 | | 0,0121 | Chitin-binding developmental regulator (IPR051940) |  | chitin binding (GO:0008061) | InterPro | Chitin binding |
| XM_059316464.1 | 50,44 | -1,28 | 0,44 | 0,000118435 | | 0,048 | Receptor Tyrosine Kinase (IPR050122) | protein phosphorylation (GO:0006468) cell migration (GO:0016477) nervous system development (GO:0007399) cell surface receptor protein tyrosine kinase signaling pathway (GO:0007169) positive regulation of kinase activity (GO:0033674) multicellular organism development (GO:0007275) | protein binding (GO:0005515) protein kinase activity (GO:0004672) ATP binding (GO:0005524) protein tyrosine kinase activity (GO:0004713) transmembrane receptor protein tyrosine kinase activity (GO:0004714) | InterPro / PANTHER | Signal transduction |
| XM_059301956.1 | 106,83 | -1,28 | 0,38 | 2,31215E-05 | | 0,0139 | Zinc-containing alcohol dehydrogenase-like protein (IPR051397) |  | zinc ion binding (GO:0008270) oxidoreductase activity (GO:0016491) | InterPro | Metabolism |
| XM_059308018.1 | 42,22 | -1,32 | 0,39 | 2,01411E-05 | | 0,0124 | Transient receptor potential channel, canonical (IPR002153) | calcium ion transmembrane transport (GO:0070588) monoatomic ion transport (GO:0006811) transmembrane transport (GO:0055085) regulation of cytosolic calcium ion concentration (GO:0051480) | calcium channel activity (GO:0005262) monoatomic ion channel activity (GO:0005216) store-operated calcium channel activity (GO:0015279) inositol 1,4,5 trisphosphate binding (GO:0070679) | InterPro / PANTHER | Ion transport |
| XR_009384131.1 | 29,91 | -1,69 | 0,45 | 7,71099E-06 | | 0,0072 |  |  |  |  | Unknown |
| XM_059315465.1 | 251,58 | -1,88 | 0,48 | 2,90931E-06 | | 0,0038 |  |  |  |  | Unknown |
| XM_059306480.1 | 32,88 | -5,62 | 1,39 | 3,89864E-07 | | 0,0008 | TVP38/Transmembrane protein 64 (IPR053069) | regulation of cytosolic calcium ion concentration (GO:0051480) |  | PANTHER | Calcium signaling |
